# Sex-Specific DNA Methylation and Gene Expression Changes in Mouse Placentas After Early Preimplantation Alcohol Exposure

**DOI:** 10.1101/2023.09.30.560198

**Authors:** Lisa-Marie Legault, Thomas Dupas, Mélanie Breton-Larrivée, Fannie Filion-Bienvenue, Anthony Lemieux, Alexandra Langford-Avelar, Serge McGraw

## Abstract

During pregnancy, exposure to alcohol represents an environmental insult capable of negatively impacting embryonic development. This influence can stem from disruption of molecular profiles, ultimately leading to manifestation of fetal alcohol spectrum disorder. Despite the central role of the placenta in proper embryonic development and successful pregnancy, studies on the placenta in a prenatal alcohol exposure and fetal alcohol spectrum disorder context are markedly lacking. Here, we employed a well-established model for preimplantation alcohol exposure, specifically targeting embryonic day 2.5, corresponding to the 8-cell stage. The exposure was administered to pregnant C57BL/6 female mice through subcutaneous injection, involving two doses of either 2.5 g/kg 50% ethanol or an equivalent volume of saline at 2-hour intervals. Morphology, DNA methylation and gene expression patterns were assessed in male and female late-gestation (E18.5) placentas. While overall placental morphology was not altered, we found a significant decrease in male ethanol-exposed embryo weights. When looking at molecular profiles, we uncovered numerous differentially methylated regions (DMRs; 991 in males; 1309 in females) and differentially expressed genes (DEGs; 1046 in males; 340 in females) in the placentas. Remarkably, only 21 DMRs and 54 DEGs were common to both sexes, which were enriched for genes involved in growth factor response pathways. Preimplantation alcohol exposure had a greater impact on imprinted genes expression in male placentas (imprinted DEGs: 18 in males; 1 in females). Finally, by using machine learning model (L1 regularization), we were able to precisely discriminate control and ethanol-exposed placentas based on their specific DNA methylation patterns. This is the first study demonstrating that preimplantation alcohol exposure alters the DNA methylation and transcriptomic profiles of late-gestation placentas in a sex-specific manner. Our findings highlight that the DNA methylation profiles of the placenta could serve as a potent predictive molecular signature for early preimplantation alcohol exposure.

**GRAPHICAL ABSTRACT:** 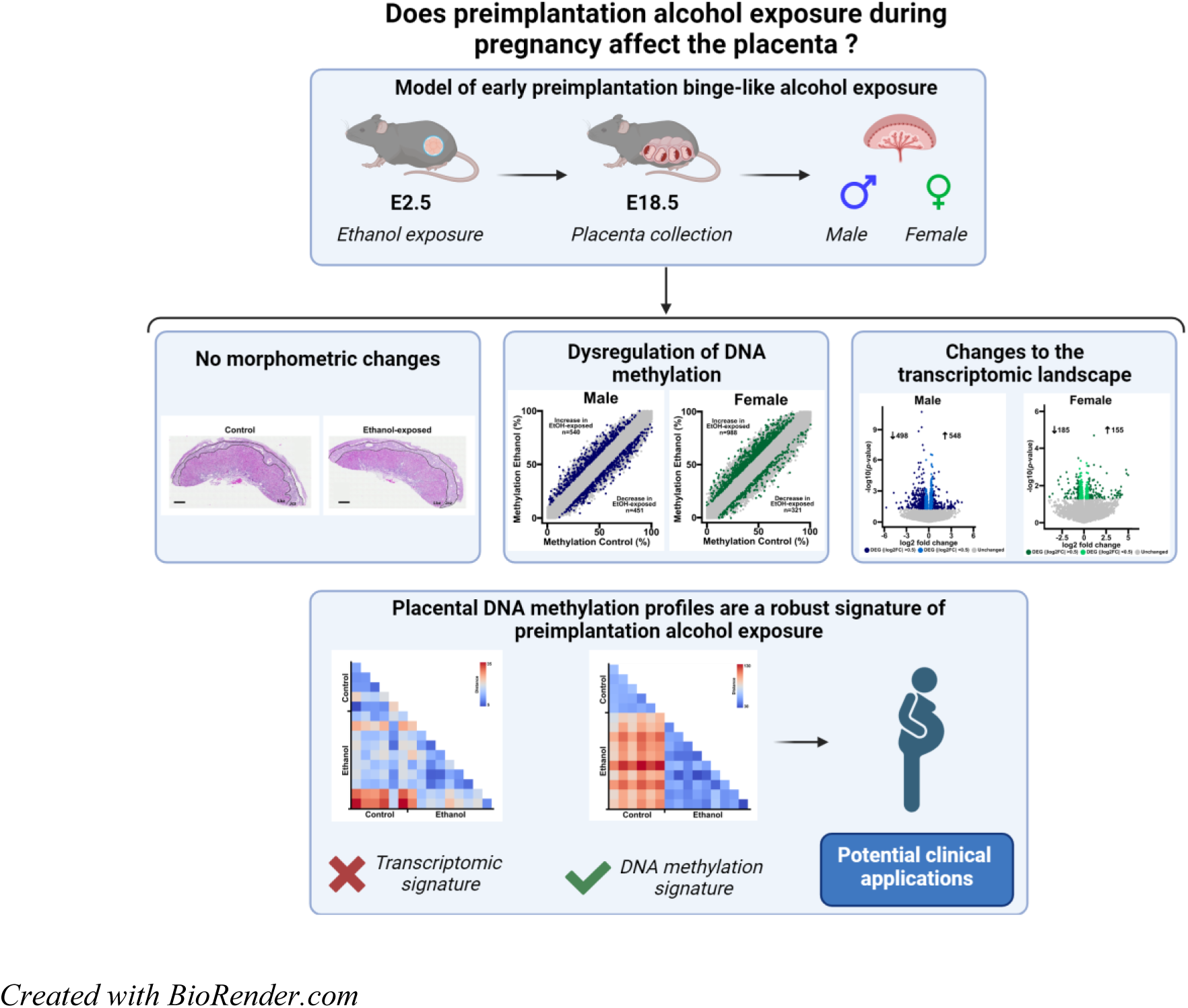

## 1. INTRODUCTION

The maternal environment is essential for a healthy pregnancy and embryonic development. Adverse effects of alcohol exposure at any point during pregnancy are referred to as fetal alcohol spectrum disorder (FASD) (Popova et al., 2023). FASD manifests as various symptoms, including cognitive impairments, learning difficulties, behavioral issues, and, in severe cases, profound intellectual disabilities and craniofacial abnormalities (Popova et al., 2023). Annually, approximately 630,000 newborns are affected by FASD worldwide (Lange et al., 2017). Its presentation and severity depend on multiple factors, including the timing, amount, and pattern of prenatal alcohol exposure (PAE) (Popova et al., 2023). Poor maternal health, exposure to other environmental insults (e.g. tobacco, drugs, contaminants), and nutritional deficiencies can also increase FASD severity (Popova et al., 2023). Currently, there is no molecular diagnostic tool for FASD, and most cases are not identified until school-aged children, when neurodevelopmental symptoms become noticeable (Popova et al., 2023). Early diagnosis and management can alleviate the impacts of FASD, while late detection often results in a need for lifelong health and social care (Popova et al., 2023).

The placenta is a transient organ that supplies oxygen, essential nutrients, hormones, and metabolites to the developing fetus. The placenta is of fetal lineage, stemming from the same pool of early embryonic blastomere cells that form the embryo, and shares its sex (Hemberger et al., 2020). During early development, the trophectoderm and inner cell mass separate into distinct lineages, and the placenta develops alongside the embryo after implantation. In mice, placental remodeling occurs as the chorion fuses with the allantois, followed by trophoblast differentiation and labyrinth structure formation (Hemberger et al., 2020). This remodeling process typically ends by mid-gestation, at E10.5 in mice and E18 in humans (Hemberger et al., 2020; Simmons et al., 2008). Proper placental development and function are crucial for the fetal growth, as the placenta regulates the availability of hormones and nutrients (Hart et al., 2019). Additionally, the placenta produces neurotransmitters, underscoring its central role in fetal brain development and highlighting a crucial placenta-brain axis (Bonnin et al., 2011; Rosenfeld, 2021). Placental abnormalities lead to a range of adverse pregnancy outcomes, including embryonic growth delays, intrauterine growth restriction, preeclampsia, and preterm birth (Coussons-Read, 2013; Wadhwa et al., 2011). Interestingly, a recent study has revealed that prenatal alcohol exposure is linked to alterations in human placental gene expression (Deyssenroth et al., 2024). This underscores the potential role of this tissue in exploring and understanding the repercussions of alcohol on embryonic development.

DNA methylation, established by DNMT3A/DNMT3B and maintained by DNMT1, is a stable modification of the DNA consisting in the covalent addition of a methyl group to cytosine (McGraw et al., 2015, 2013; Shaffer et al., 2015). By regulating gene expression, DNA methylation plays a crucial role in numerous biological pathways, thereby contributing significantly to fetal development. Importantly, DNA methylation seems to act like a sensor to exposure to environmental contaminants and/or infections (Breton-Larrivée et al., 2019; Maurice et al., 2021; Pierre et al., 2020). Changes in DNA methylation patterns in the placenta have been described as associated with fetal growth restriction (Banister et al., 2011; Lee et al., 2021). Strikingly, recent studies have highlighted that alteration in placental DNA methylation patterns in response to harmful environmental exposures (e.g., maternal smoking, prenatal air pollution) can potentially affects fetal growth (Broséus et al., 2024; Everson et al., 2021). These findings underscore the importance of assessing the impact of prenatal alcohol exposure on placental DNA methylation patterns to gain a better understanding of the holistic pathways by which prenatal alcohol exposure affects fetal development.

The preimplantation period is a critical phase in embryonic development, in which the parental DNA methylation marks are erased and the epigenome is remodeled to initiate the developmental program (Canovas and Ross, 2016; Shaffer et al., 2015). Since pregnancy is typically detectable only after implantation, when human chorionic gonadotropin hormone production begins, there is a 6-day window of high risk for PAE during the preimplantation development stage (Cole, 2009). Given the increasing number of women of 18–34 years old consuming alcohol, including binge drinking (World Health Organization, 2018), and the substantial number of unintended pregnancies worldwide (approximately 40%; totaling 85 million annually) (Sedgh et al., 2014), many women may unknowingly expose their developing embryos to high levels of alcohol during the initial weeks of pregnancy. Thus, it is essential to understand the impact of alcohol during this critical phase and more particularly a single binge-like prenatal alcohol exposure which can lead to more severe brain defects and cognitive/behavioral outcomes compared to chronic exposure, due to its ability to produce the highest peak blood alcohol concentration (Maier and West, 2001). Despite its importance for proper embryonic development and its heightened susceptibility to adverse environmental exposures, the preimplantation period has been understudied in relation to PAE, with only a few models focusing exclusively on this early developmental timepoint (Breton-Larrivée et al., 2023; Haycock and Ramsay, 2009; Legault et al., 2021, 2018).

In our recent study using a preclinical mouse model of early preimplantation binge-like alcohol exposure, we reported sex-specific DNA methylation changes in the forebrains of embryos at mid-gestation (E10.5) (Legault et al., 2021). We also observed an increased incidence of morphological defects, including growth delays, in embryos exposed to ethanol at E10.5 and E18.5 (Breton-Larrivée et al., 2023; Legault et al., 2021). These disruptions could be potentially imputed to changes in DNA methylation at imprinting control regions (ICRs) which are associated with fetal growth restriction (Bourque et al., 2010; O. Koukoura et al., 2011; Ourania Koukoura et al., 2011). Despite all this evidence, a significant knowledge gap remains regarding the effects of early preimplantation alcohol exposure on the placenta. The aim of the study was to define impacts of early preimplantation prenatal alcohol exposure on the late-gestation placenta using our preclinical mouse model.

## 2. MATERIAL AND METHODS

### 2.1. Mouse model of preimplantation alcohol exposure

Eight-week-old female and male C57BL/6 mice (Charles River Laboratories) were mated, and pregnancy with E0.5 embryos was confirmed by the presence of a visible copulatory plug the following morning. Pregnant females were separated from the males and housed together (*n=*2–3) in a 12-h light/dark cycle with unrestricted access to food and water. Using a recognized prenatal binge-like alcohol exposure paradigm (Haycock and Ramsay, 2009; Kleiber et al., 2013; Mantha et al., 2014; Padmanabhan and Hameed, 1988) and as previously described (Breton-Larrivée et al., 2023; Legault et al., 2021), pregnant females were subcutaneously injected with two doses of either 2.5 g/kg of 50% ethanol or an equivalent volume of saline at 2-h intervals on E2.5 (the 8-cell stage), then housed under similar conditions with limited handling during gestation.

### 2.2. Tissue collection and analysis

At E18.5, pregnant females were euthanized and embryos and placentas were collected. The maternal tissue layer was then removed from the placenta. The embryos and placentas were weighed, and the placental area was calculated using a Leica stereo microscope and Leica LasX software (v1.4.4). Placentas were flash frozen in liquid nitrogen and stored at -80°C for subsequent DNA and RNA extraction. The sex of each embryo and its placenta was determined by qPCR for *Ddx3x* and *Ddx3y* (LightCycler 96, Roche) (Breton-Larrivée et al., 2023; Legault et al., 2021).

### 2.3. Histological analysis

On E18.5, placentas were collected with the maternal decidua to preserve structural integrity. Placentas were fixed in 4% paraformaldehyde, post-fixed in 70% ethanol for 48 h, and then divided into two identical sections for paraffin embedding. Cross-sections (5 µm thick) were stained with hematoxylin and eosin and imaged on a Zeiss Zen Axioscan Slide Scanner (Legault et al., 2021). Image processing and junctional and labyrinth zone measurements were performed with ImageJ (v1.52).

### 2.4. DNA/RNA extraction and library preparation

To capture the spectrum of alterations in response to ethanol exposure (Breton-Larrivée et al., 2023; Legault et al., 2021), and to minimize potential bias associated with a small sample size, we use a larger number of animals in the ethanol-exposed group compared to the controls. We randomly selected 6 control placentas (1 male and 1 female per litter from 3 different litters) and 10 ethanol-exposed placentas (1 male and 1 female per litter from 5 different litters) with normal morphometric measurements and no visible morphological defects. The corresponding embryos also developed normally. Whole placentas were ground in liquid nitrogen and equivalent amounts of each sample were used for DNA and RNA extraction using QIAamp DNA Micro (Qiagen #56304) and RNeasy Mini (Qiagen #74004) kits, respectively, following the manufacturer’s recommendations. DNA and RNA were quantified on a QuBit Fluorimeter using Broad-Range DNA assay (ThermoFisher #Q32853) and High Sensitivity RNA assay (ThermoFisher #Q32852) kits, respectively, following the manufacturer’s recommendations.

DNA (1 µg) was used to produce Methyl-Seq libraries using a SureSelectXT Methyl-Seq Target Enrichment System with a mouse enrichment panel (Agilent #G9651B and #5191-6704) following the manufacturer’s recommendations. After final amplification/indexation, libraries were quantified using a QuBit fluorimeter with a High Sensitivity DNA assay kit (ThermoFisher #Q32854) and quality was assessed using a BioAnalyzer before paired-end sequencing on a NovaSeq6000 S4 at the Genome Quebec core facility. We obtained 112–139 M reads per sequenced library.

High-quality RNA (500 ng) was used to produce mRNA-Seq libraries using NEBNext mRNA stranded library kits at the Genome Quebec core facility, followed by paired-end sequencing on a NovaSeq6000 S4. We obtained 26–43 M reads per sequenced library.

### 2.5. Bioinformatics analysis

Methyl-Seq results were analyzed using the GenPipes Methyl-Seq pipeline (v3.3.0) (Bourgey et al., 2019). Reads were aligned to the mouse reference genome (mm10) and methylation calls were obtained with Bismark (v0.18.1) (Krueger and Andrews, 2011). The R package MethylKit (version 1.8.1) (Akalin et al., 2012) was used to obtain differentially methylated regions (DMRs) based on the Benjamini–Hochberg false discovery rate procedure. We used fixed parameters, such as 100 base pair (bp) stepwise tiling windows and a threshold of *q* < 0.01 and ≥10% difference of methylation between conditions. DNA methylation levels were calculated as the average methylation of all CpGs within a tile for all the samples within a condition (minimum 3 samples/condition/sex at ≥ 10X sequencing depth) (Legault et al., 2021, 2020). The bisulfite conversion rate (> 96%) and number of CpGs per tile were obtained using a custom Perl script (Legault et al., 2021).

Annotation of all analyzed tiles was conducted using Homer (version 4.10.1) with the mouse mm10 reference genome (Heinz et al., 2010). Functional enrichment analyses were performed with Metascape (Zhou et al., 2019) using differentially methylated tiles located in genes as an input. We obtained CpG island coordinates of the mm10 genome from the UCSC Genome Browser (Karolchik et al., 2004), then delineated CpG shores and shelves by extending 0–2 and 2–4 kb from them, respectively, as described previously (Legault et al., 2021, 2020).

For mRNA-Seq data, post-sequencing bioinformatics analyses were performed on the mouse reference genome (mm10) using the GenPipes RNA-Seq pipeline (v4.1.2) (Bourgey et al., 2019), including tools such as Trimmomatic (v0.36) (Bolger et al., 2014), STAR (v2.5.3) (Dobin et al., 2013), Picard (v2.9.0), and BWA (v0.7.12) (Li and Durbin, 2009). DEGs were obtained using the R package DESeq2 (v1.24.0) (Love et al., 2014) including multiple testing correction using the Benjamini-Hochberg false discovery rate procedure. Genes with *p*-values ˂ 0.05 were considered significant.

For the deconvolution analysis, we formatted the Single Nuclei data of E18.5 mouse placenta from a relevant preprint article (Fu et al., 2024) to produce an Atlas usable to deconvolute our bulk RNA-seq dataset. The final Single Nuclei dataset consisted of 9779 cells across 28 cell populations. Firstly, we normalized the data to 1000 counts per cell by the scanpy.pp.normalize_per_cell function followed by a log transformation with the scanpy.pp.log1p function. Then we selected the top 4000 highly variable genes and calculated the centroids of cell types to lauch the AutoGeneS (v1.0.4) (Aliee and Theis, 2021) initialization function. The optimize function of AutoGeneS was then used with 5000 generations for 400 genes and the deconvolution was finally run on our raw counts bulk RNA-seq matrix with the NuSVR model. To further investigate if a particular cell type was enriched or diminished following exposure, we took the mean value associated with each cell composition and compared the means (Ctrl vs EtOH) using a non-parametric test (Mann-Whitney-Wilcoxon) and adjusted the p-values for multiple testing with the FDR correction.

To establish a molecular signature of early preimplantation PAE, we used L1-regularized linear regression (LASSO) (Sohn et al., 2009; Xu et al., 2022) with cross-validation for the gene expression and DNA methylation profiles. The curse of dimensionality (*i.e.*, challenges stemming from training models on datasets with a large number of features relative to samples) was addressed using the Synthetic Minority Over-sampling Technique to counteract sample imbalances (Taft et al., 2009). A random forest classifier was trained on these balanced datasets to emphasize the significance of the selected DEGs and DMRs in the classification. The specific regions used to establish the signature were among the top ranked based on feature importance. The efficacy of the identified DEGs or DMRs to differentiate the two groups was validated by two-sample *t-*test and Wilcoxon rank-sum test, while ROC curves were used to assess the discriminatory capacity of each DEGs or DMRs (Sohn et al., 2009).

### 2.6. Statistics

Statistical analyses not included in bioinformatic pipelines were performed using R (version 3.5.0) or GraphPad Prism (version 9.5.0). Statistical significance was calculated using unpaired *t*-tests with Welch’s correction and an *F* test for variance or two-proportion *z*-test. All data represent the mean ± standard deviation (SD). *p*-values ˂ 0.05 were considered significant.

## 3. RESULTS

### 3.1. Early preimplantation alcohol exposure does not induce morphometric changes in the placenta but induces lower embryo weight in males

In this study, we examined the effect of early preimplantation binge alcohol exposure on late placental development using an established FASD mouse model (Breton-Larrivée et al., 2023; Legault et al., 2021). Briefly, to simulate short-term exposure to a high level of alcohol, pregnant females were given two doses (2.5 g/kg) of ethanol at 2-hour intervals during preimplantation (at the 8-cell stage (E2.5)). We previously showed that this increases the average blood alcohol level to 158.31 mg/dL (Legault et al., 2021). At late gestation (E18.5), we collected the embryos and placentas to assess the overall impacts of alcohol on gestational outcomes, focusing first on basic morphometric measurements. All measurements were compared within the same sex.

The proportion of male embryos is unchanged between control (47/86 – 55%) and ethanol-exposed group (62/101 – 61%) (Fig. 1A). Additionally, there were no significant effects of early preimplantation alcohol exposure on litter size, the number of males and females per litter (Fig. 1B), or the placental area and weight (Fig. 1C, 1D). Weight of ethanol-exposed male embryos is significantly decreased compared to controls (1.134g ± 0.067 vs 1.195g ± 0.102, respectively; *p*<0.01), while no difference was observed in females (Fig. 1E). Despite the reduced weights of male embryos, the placental efficiency (*i.e*., the embryo-to-placental weight ratio) was similar between control and ethanol-exposed embryos for both sexes (Fig. 1F).

**Figure 1:**
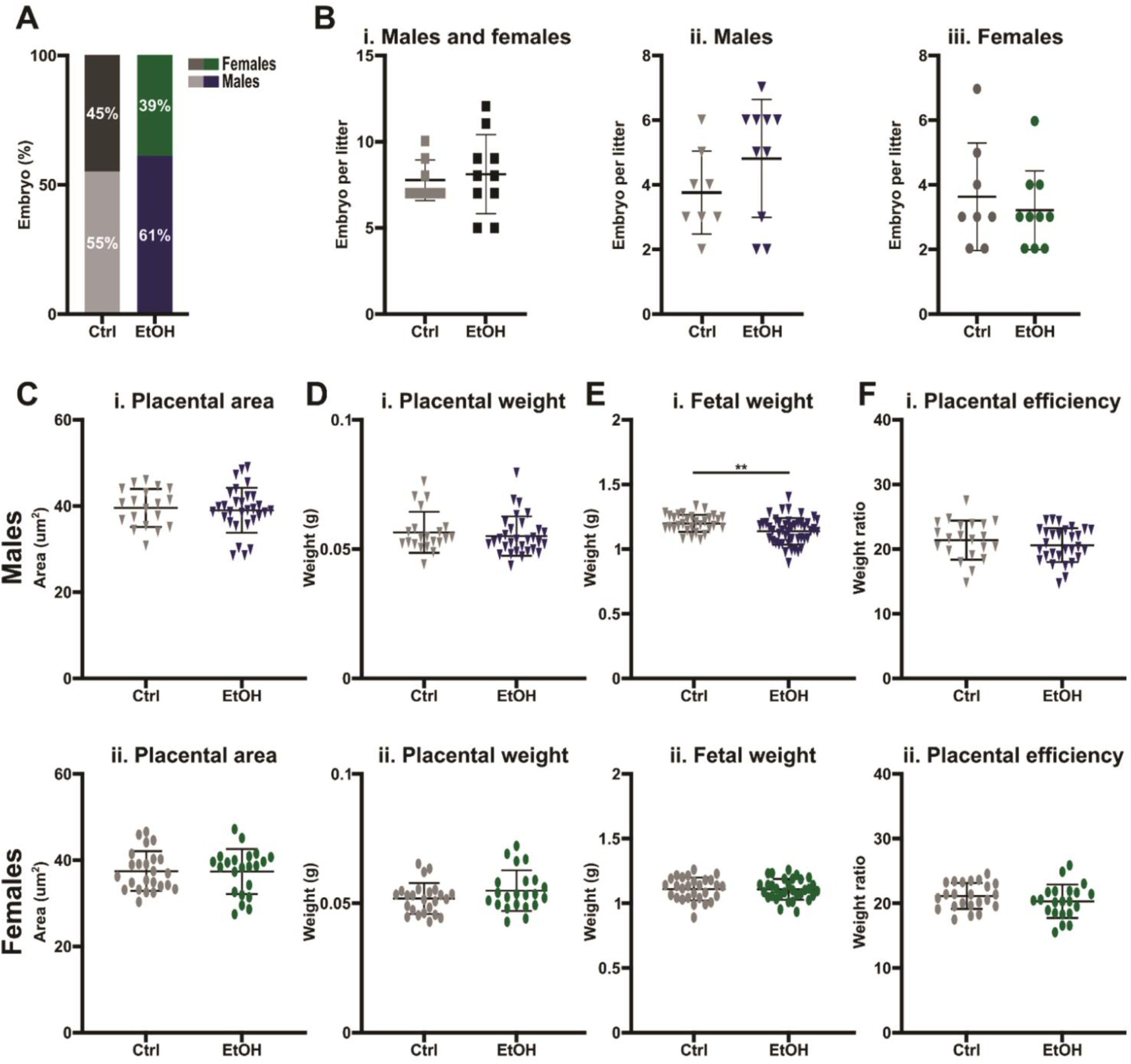
Absence of morphometric alterations in E18.5 placentas following preimplantation alcohol exposure. Evaluation of the impact of preimplantation alcohol exposure on **A)** sex ratios of litters (control: males *n*=47, females *n*=39; ethanol-exposed: males *n*=62, females *n*=39); **B) i)** total embryos, number of **ii)** male and **iii)** female embryos per litter (control: *n*=8 litters; ethanol-exposed: *n*=10 litters); **C)** placental surface area in **i)** males and **ii)** females (control embryos: males, *n*=19, females, *n*=24; ethanol-exposed embryos: males *n*=30, females *n*=22; 6 litters, excluded litters used for placentas with decidua); **D)** placental weight in **i)** males and **ii)** females (control embryos: males, *n*=19, females, *n*=24; ethanol-exposed embryos: males, *n*=30, females, *n*=22; 6 litters, excluded litters used for placentas with decidua); **E)** embryo weight in **i)** males and **ii)** females (control embryos: males, *n*=31, 8 litters; females, *n*=28, 8 litters; ethanol-exposed embryos: males, n=46, 10 litters; females, *n*=32, 8 litters); **F)** placental efficiency (embryo-to-placental weight ratio) in **i)** males and **ii)** females (control embryos: males, *n*=19, females, *n*=24; ethanol-exposed embryos: males, *n*=30, females, *n*=22; 6 litters, excluded litters used for placentas with decidua). All data represent the mean ± standard deviation (SD). Significant differences were assessed by *t*-test with Welch’s correction. **: *p*<0.01. Ctrl: control, EtOH: ethanol-exposed.

When investigating the impact of early alcohol exposure on the overall composition of placental cells, we observed no significant differences in the sizes or thicknesses of the junctional and labyrinth zones between the groups. Additionally, the ratios of their respective areas/thicknesses to the total area/thickness showed no significant differences (Fig. S1). However, in female ethanol-exposed placentas, the junctional zone thickness, and the ratio of the junctional zone area to the total area presented significantly increased variance compared to control (*p*˂0.05) (Fig. S1).

Overall, these findings suggest that early preimplantation alcohol exposure does not significantly impact placental morphology during late gestation but does have a developmental effect on male embryos.

### 3.2. Sex-specific DNA methylation dysregulation in late-gestation placentas following early preimplantation alcohol exposure

To examine the impact of preimplantation alcohol exposure on the DNA methylation patterns of late-gestation placentas, we performed genome-wide DNA methylation analyses on control and ethanol-exposed placentas. After establishing sex-specific DNA methylation profiles in E18.5 placentas, we compared the average DNA methylation levels between control and ethanol-exposed male or female placentas, and identified 751,083 tiles (100bp) with sufficient coverage in both data sets, allowing us to investigate shared and sex-specific DNA methylation differences induced by early preimplantation alcohol exposure (Fig. S2A).

Strikingly, we identified 991 and 1309 differentially methylated regions (DMRs; ≥10% difference between conditions) in response to early preimplantation alcohol exposure that were exclusive to males and females (Fig. 2A, 2C; Table 1 and S1), respectively, while only 21 DMRs were shared between the sexes (Fig. S2A; Table 2 and S1). Male placentas had nearly equal distributions of DMRs, displaying both increased (540 – 54%) and decreased (451 – 46%) methylation after ethanol exposure (Fig. 2A). In contrast, DMRs in female placentas were predominantly increased with ethanol exposure (988 – 75% vs. 321 – 25%) (Fig. 2C). Most DMRs were annotated as introns (male, 41%; female, 42%) or intergenic regions (male, 39%; female, 37%) (Fig. 2B, 2D). We observed larger DNA methylation average changes between control and ethanol-exposed placentas in DMRs located in non-coding regions (controls 44% vs ethanol-exposed 51%) and in exons (controls 47% vs ethanol-exposed 52%) in males, while in females, larger average changes were observed for DMRs located in 3′ UTRs (controls 53% vs ethanol-exposed 63%) and introns (controls 45% vs ethanol-exposed 54%) (Fig. 2B, 2D).

**Figure 2:**
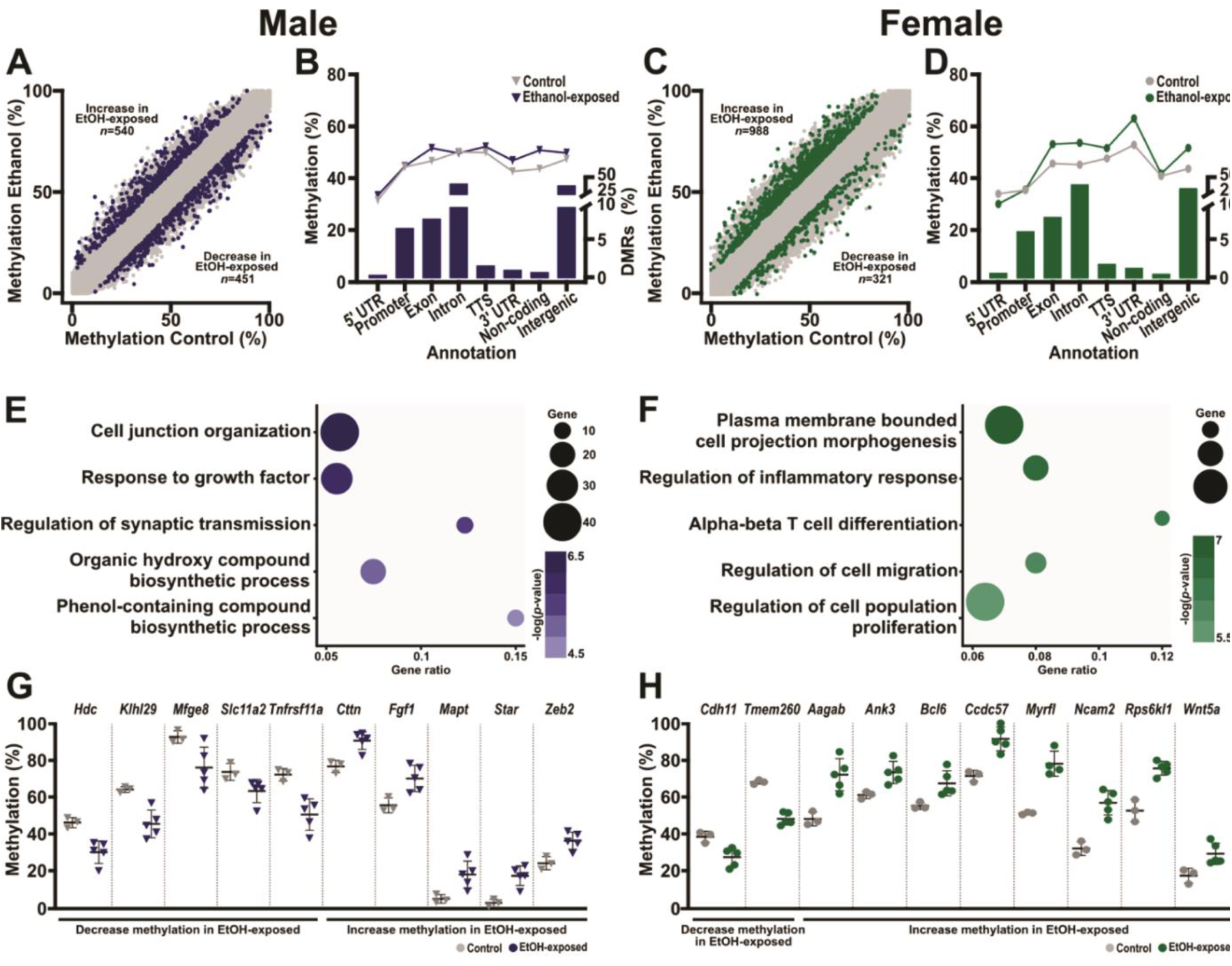
Sex-specific DNA methylation dysregulation in the placenta following early preimplantation alcohol exposure. Scatterplot of differentially methylated regions (DMRs) between control and ethanol-exposed **A)** male and **C)** female placentas. Blue (male) or green (female) dots represent tiles with ≥ 10% methylation changes in ethanol-exposed placentas compared to controls (number of DMRs: male: 991, female: 1309). CpG methylation levels in control and ethanol-exposed **B)** male and **D)** female placentas based on DMR annotations (left Y-axis) and DMR distributions across genomic annotations (right Y-axis). Top enriched pathways in **E)** male (542 unique gene DMRs) and **F)** female (749 unique gene DMRs) DMRs based on Metascape analysis. CpG methylation levels of top changed DMRs or DMRs associated with the top enriched pathways shown in **E** and **F** in control and ethanol-exposed **G)** male (left; grey/blue) and **H)** female placentas.

**Table 1:**
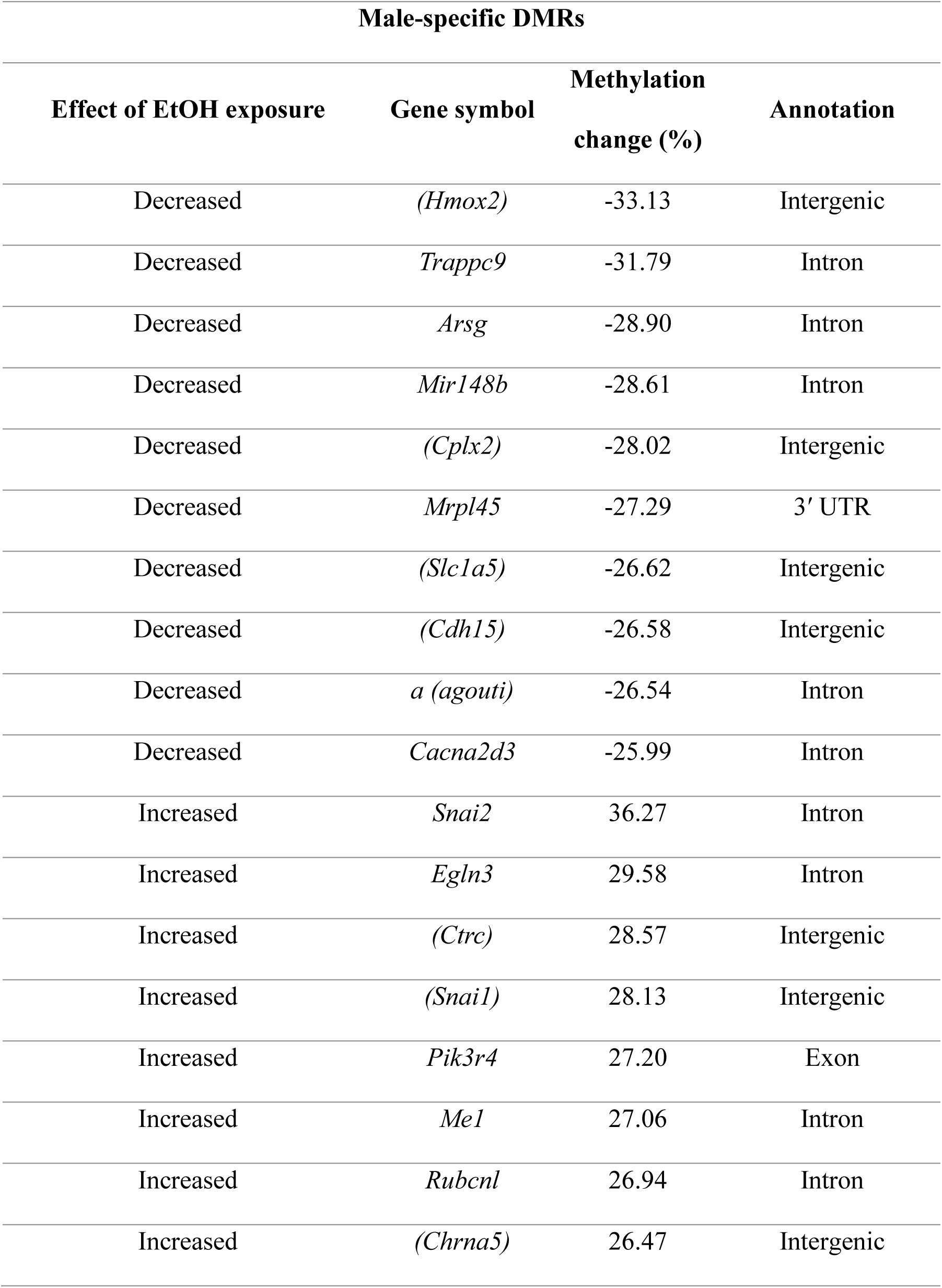
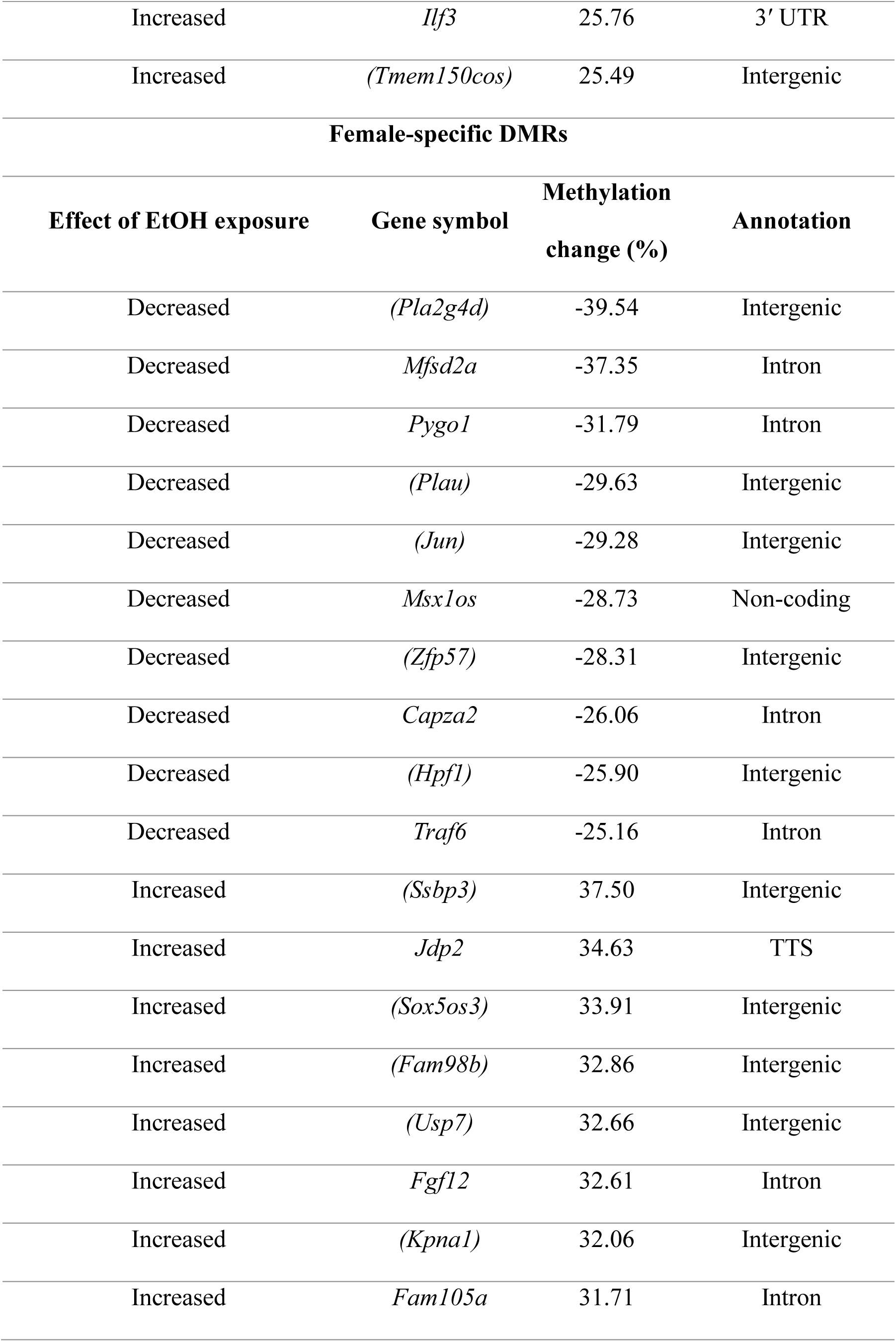
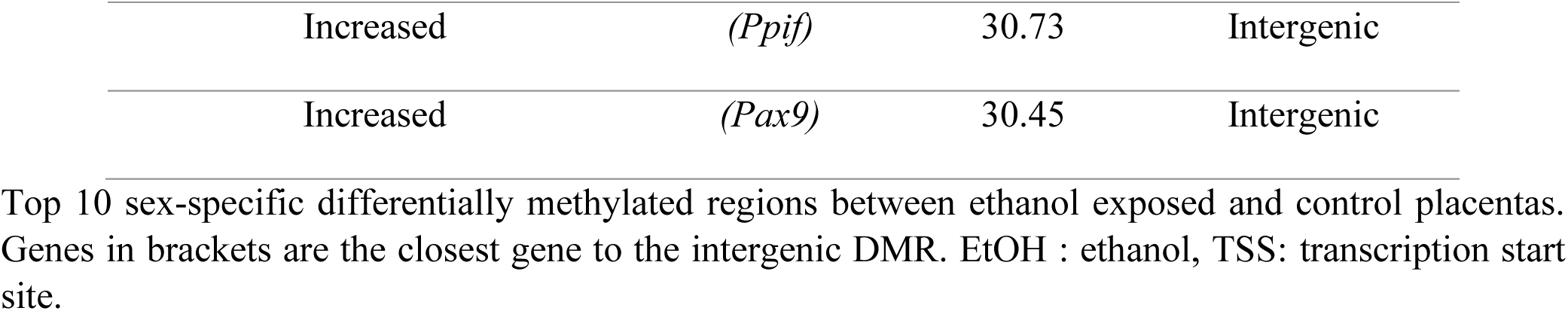
Top 10 sex-specific DMRs.

**Table 2:** Top 10 shared DMRs between male and female placentas.

| Shared top DMRs |  |  |  |  |  |
| --- | --- | --- | --- | --- | --- |
| Effect of EtOH<br>(males) | Gene<br>symbol | Methylation<br>change (%)<br>(males) | Effect of EtOH<br>(females) | Methylation<br>change (%)<br>(females) | Annotation |
| *Cdk8* | Decreased | -23.33 | Decreased | -12.21 | Intron |
| *(Mirt1)* | Decreased | -13.04 | Decreased | -12.24 | Intergenic |
| *Hs3st2* | Decreased | -12.54 | Decreased | -10.83 | Exon |
| *Tmem267* | Decreased | -12.51 | Decreased | -10.67 | Promoter-TSS |
| *Rbm3os* | Decreased | -10.07 | Increased | 15.50 | Promoter-TSS |
| *(Ifngas1)* | Increased | 15.22 | Decreased | -20.68 | Intergenic |
| *(Slc22a19)* | Increased | 10.02 | Decreased | -10.61 | Intergenic |
| *Tro* | Increased | 10.12 | Increased | 13.77 | Promoter-TSS |
| *Galp* | Increased | 15.12 | Increased | 10.21 | Exon |
| *(Spry1)* | Increased | 15.01 | Increased | 15.70 | Intergenic |
| *Clstn3* | Increased | 13.36 | Increased | 15.06 | Exon |
| *(Spry1)* | Increased | 13.08 | Increased | 12.83 | Intergenic |
| *Spry1* | Increased | 13.00 | Increased | 16.15 | Intergenic |
| *(Spry1)* | Increased | 12.05 | Increased | 11.63 | Intergenic |
| *Socs5* | Increased | 11.55 | Increased | 13.67 | Intron |
| *Psd3* | Increased | 11.40 | Increased | 10.02 | Intron |
| *Socs5* | Increased | 11.02 | Increased | 11.20 | Exon |
| *Wwox* | Increased | 10.80 | Increased | 10.94 | Intron |
| *Trh* | Increased | 10.18 | Increased | 11.33 | Intergenic |

Top 10 differentially methylated regions shared by males and females between alcohol-exposed and control placentas. Genes in brackets are the closest gene to the intergenic DMR. EtOH : ethanol, TSS: transcription start site.

A detailed examination of the distributions of sex-specific DMRs across various CpG methylation levels revealed a consistent pattern between the sexes. DNA methylation tends to increase in genomic regions with higher basal methylation levels (50-100%), while it decreases in genomic regions with lower basal methylation levels (0-50%), as shown by the rightward shift of the EtOH-exposed associated curve (Fig. S3A, S3C). This trend was especially pronounced for female-specific DMRs (Fig. S3C). Although some sex-specific DMRs displayed methylation changes exceeding 20%, the majority exhibited changes in the range of 10 to 15% (Fig. S3B, S3D).

To identify pathways associated with sex-specific DMRs in ethanol-exposed placentas, we performed gene annotation enrichment analyses. First, we removed DMRs associated with intergenic regions. For males, the remaining 607 DMRs represented 542 unique genes. Enriched pathways included cell junction assembly (*Cttn*, *Mapt*), response to growth factors (*Fgf1*, *Zeb2*), negative regulation of synaptic transmission (*Cttn*, *Mapt*), organic hydroxy compound biosynthesis, (*Slc11a2*, *Star*), and phenol-containing compound biosynthesis (*Hdc*, *Nr4a2*) (Fig. 2E,2G). For females, the remaining 821 DMRs corresponded to 749 unique genes. The top enriched pathways differed from in males, including plasma membrane-bounded cell projection morphogenesis (*Ank3*, *Cdh11*), regulation of inflammatory responses (*Bcl6*, *Foxj1*), alpha-beta T cell differentiation (*Bcl6*, *Foxj1*), negative regulation of cell migration (*Rbp4*) and negative regulation of cell population proliferation (*Foxj1*, *Wnt5a*) (Fig. 2F,2H).

Interestingly, 1 genic male-DMRs (*Rapgef2*) and 5 genic female-DMRs (*Hrh2*, *Kmt2a*, *Large1*, *Serpine1*, *Trim58*) were associated with genes differentially expressed (n=40) in the placentas of women who reported heavy drinking during pregnancy (Deyssenroth et al., 2024). Among these, *Rapgef2*, involved in calcium ion binding and signal transducer activity, has been associated with cognitive impairment (Jang et al., 2021).

These findings reveal that early preimplantation alcohol exposure results in sex-specific DNA methylation dysregulation in late-gestation placentas, with females showing a higher susceptibility. Furthermore, these sex-specific alterations in DNA methylation are linked to specific biological pathways.

### 3.3. Female ethanol-exposed placentas tend to lose methylation in CpG-rich regions

To determine whether the local genomic sequence contributes to the inherent sex-specific susceptibility of DMRs, we examined their CpG contexts in relation to CpG islands (CGIs). Relative to control, most male and female ethanol-exposed placentas showing decreased and increased DMRs were located in open sea regions, far from CGIs (Fig. 3A). Regarding the DMRs situated in CGIs, in males, we predominantly observed DMRs with a gain of methylation (68 increased vs. 11 decreased). Conversely, in females, a larger proportion of DMRs with methylation loss was observed (45 decreased vs. 37 increased) (Fig. 3A). For both sexes, the largest changes in methylation levels occurred in open sea regions and shores, and the smallest changes mainly occurred in CGIs (Fig. 3B).

**Figure 3:**
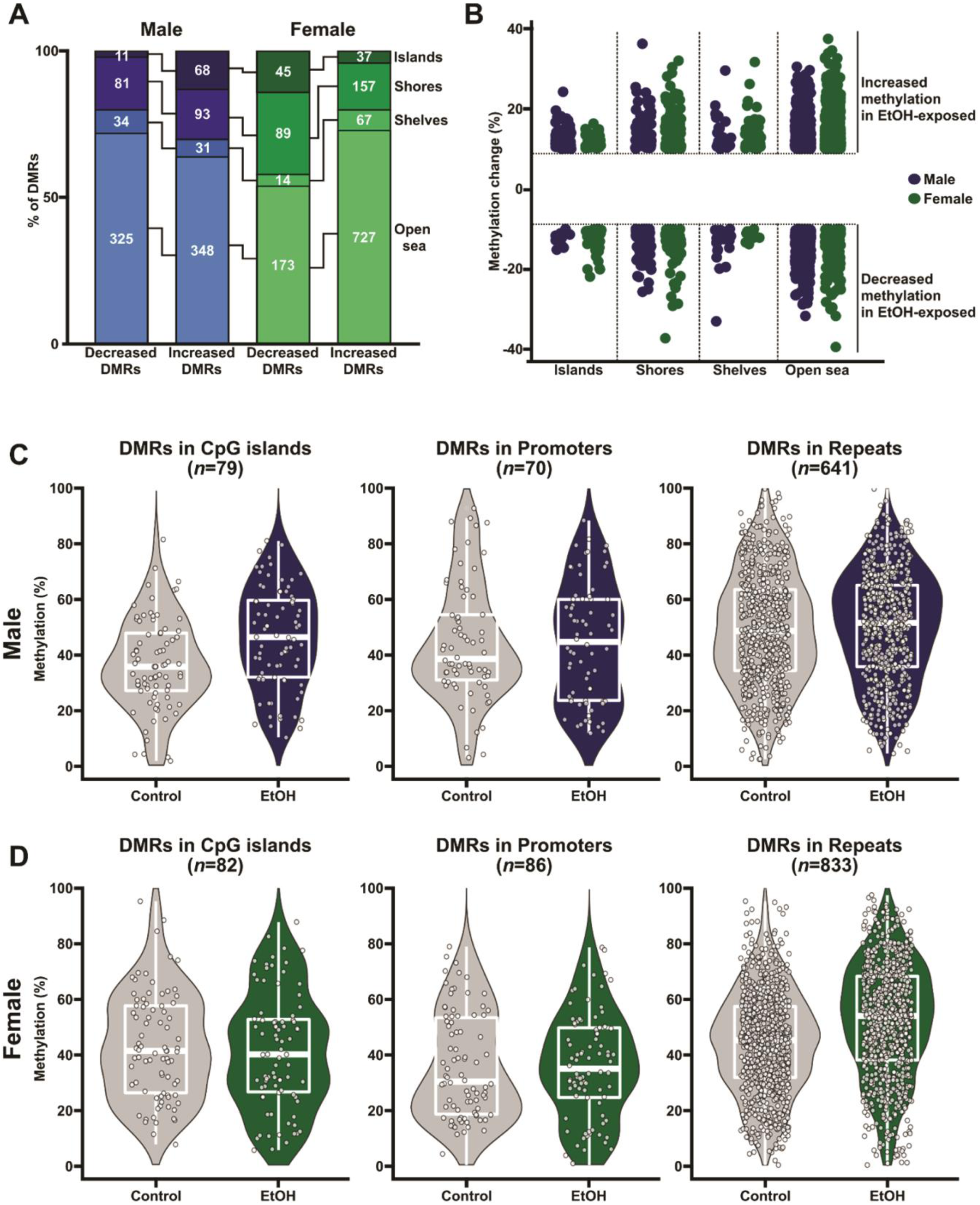
Female ethanol-exposed placentas tend to lose methylation in CpG rich regions. **A)** Distribution of DMRs between control and ethanol-exposed male (blue) and female (green) placentas based on their proximity to CpG-rich regions (CpG islands (CGIs). Regions are defined as shores (≤ 2 kb from CGIs), shelves (2–4 kb from CGIs), and open sea (≥ 4 kb from CGIs). **B)** Methylation differences in male- (blue) and female- (green) specific DMRs based on their proximities to CGIs. DNA methylation distributions in different CpG density contexts in **C)** males and **D)** females control and ethanol-exposed placentas. Ctrl: control, EtOH: ethanol-exposed.

For a more comprehensive assessment of methylation differences in sex-specific DMRs based on their CpG contexts, we compared the methylation profiles of DMRs in regions of different CpG density, notably CGIs, promoters, and repeat regions (Fig. 3C, 3D). Generally, male DMRs in CGIs had higher methylation in ethanol-exposed placentas than in controls. This difference lessened with decreasing CpG density, consistent with the fact that DMRs in repeat regions displayed both decreased and increased methylation levels in male ethanol-exposed placentas compared to controls (Fig. 3C). Interestingly, we observed the opposite trend when examining the methylation profiles of female DMRs across different annotations. While we observed generally lower methylation levels for DMRs in CGIs in ethanol-exposed placentas, the methylation levels of DMRs in promoters and particularly in repeat regions increased in female ethanol-exposed placentas compared to controls (Fig. 3D).

Taken together, our findings suggest that early preimplantation alcohol exposure differentially impacts the placental epigenome depending on sex and CpG density.

### 3.4. DMRs shared by male and female late-gestation placentas after early preimplantation alcohol exposure are related to growth and development

Although our primary focus was to identify sex-specific DMRs in the placenta resulting from early preimplantation alcohol exposure, we also observed shared affected regions between sexes. Of the 21 shared DMRs (Table 2), 17 (80%) DMRs increased or decreased consistently in both sexes, while 4 (20%) displayed opposing changes (Fig. S4A). Regions of interest included DMRs located in genes such as *Galp* (neuropeptide signaling), *Psd3* (protein transduction), or *Wwox* (apoptotic signaling) (Fig. S4B). A total of 12 (57%) DMRs were located in promoters or intragenic regions. Pathway enrichment analysis of these regions revealed enrichment in growth regulators (Fig. S4C, S4D).

For a broader understanding of how preimplantation alcohol exposure alters the DNA methylation patterns of specific genes in both male and female placentas, we performed DMRs comparisons based on gene name annotations instead of specific genomic tile-positions. After excluding intergenic DMRs, we compared a total of 607 male-specific DMRs (542 genes) to 821 female-specific DMRs (749 genes) located in promoters and intragenic regions. Along with the 21 shared DMRs previously described, we uncovered 69 shared genes across 89 DMRs with DNA methylation differences in male or female ethanol-exposed placentas (Fig. S5A). These included methylation changes that were consistent (39) and opposing (50) between the sexes (Fig. S5A, S5B). Pathway enrichment analysis revealed pathways related to the reproductive process (*Bmp6*, *Prdm14*, *Runx1*), cell morphogenesis (*Prdm14*, *Ptprk*), transmembrane transport activity, myeloid cell development, and response to growth factors (*Bmp6*, *Prdm14*, *Ptprk*, *Runx1*), (Fig. S5C, S5D).

These findings indicate that the dysregulated pathways shared between male and female placentas in response to preimplantation alcohol exposure could contribute to pathogenicity of diverse symptoms including growth defects.

### 3.5. Early preimplantation alcohol exposure leads to sex-specific gene expression changes in late-gestation placentas

To further our understanding of the molecular impact of preimplantation alcohol exposure on the late-gestation placenta, we performed gene expression profiling using mRNA-Seq of the same set of samples used for DNA methylation analysis. We used a significance threshold of *p*<0.05 to identify differentially expressed genes (DEGs) between ethanol-exposed and control placentas. We identified 1046 male-specific DEGs, 340 female-specific DEGs, and 54 shared DEGs (Fig. 4A-D; Fig. S6A; Table 3 and 4). In males, 498 genes (48%) showed decreased expression in ethanol-exposed placentas, while 548 genes (52%) showed increased expression (Fig. 4A, 4C). Of the 340 female-specific DEGs in ethanol-exposed placentas, 185 (54%) and 155 (46%) showed decreased and increased expression relative to the control, respectively (Fig. 4B, 4D).

**Figure 4:**
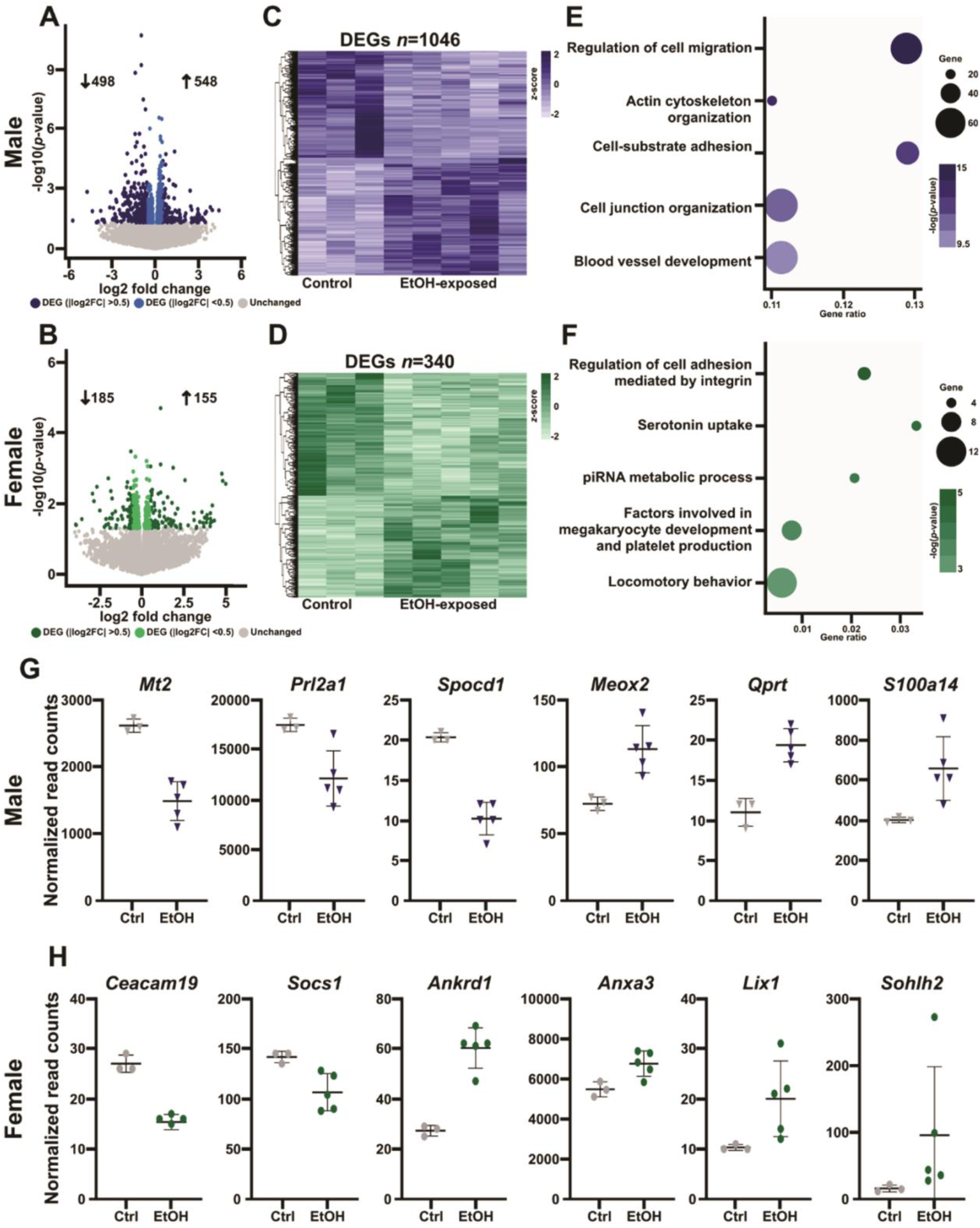
Gene expression profiles are especially affected in the male placenta following preimplantation alcohol exposure. **A-B)** Differential expression analyses between control and ethanol-exposed placenta in **A)** males and **B)** females. Colored dots represent statistically significant DEGs (*p*<0.05; males, 1046; females, 340). Darker dots represent DEGs with log2 fold change values > |0.5| (males, *n*=359; females, *n*=157). **C-D)** Heatmaps of DEG expression levels (*z*-scores) in control and ethanol-exposed **C)** male and **D)** female placentas. **E-F)** Top pathways for **E)** male (1046) and **F)** female (340) DEGs based on Metascape analysis. **G-H)** Normalized read counts of selected DEGs in **G)** male and **H)** female ethanol-exposed placentas. Data represent the mean ± SD. Ctrl: control, EtOH: ethanol-exposed.

**Table 3:**
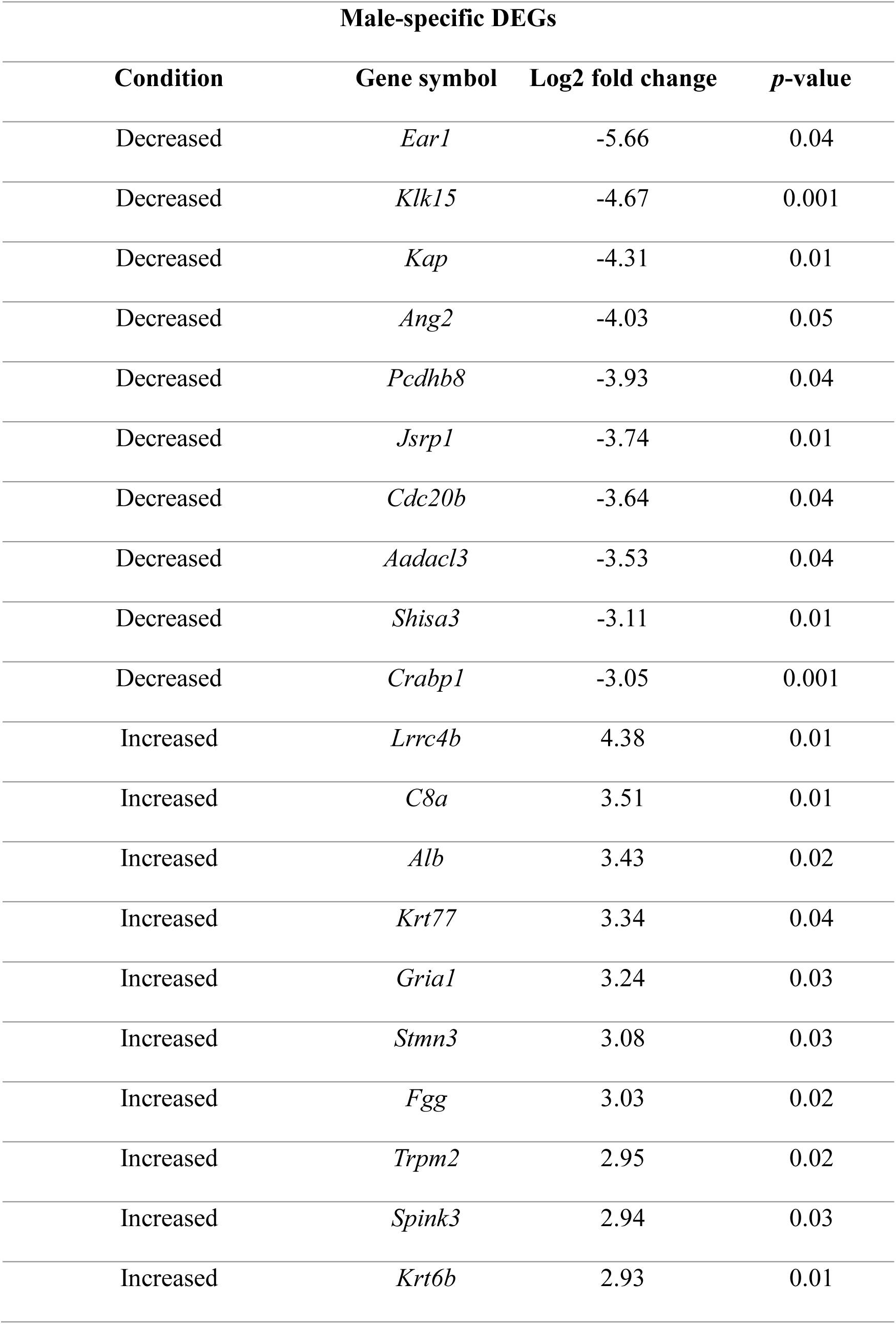
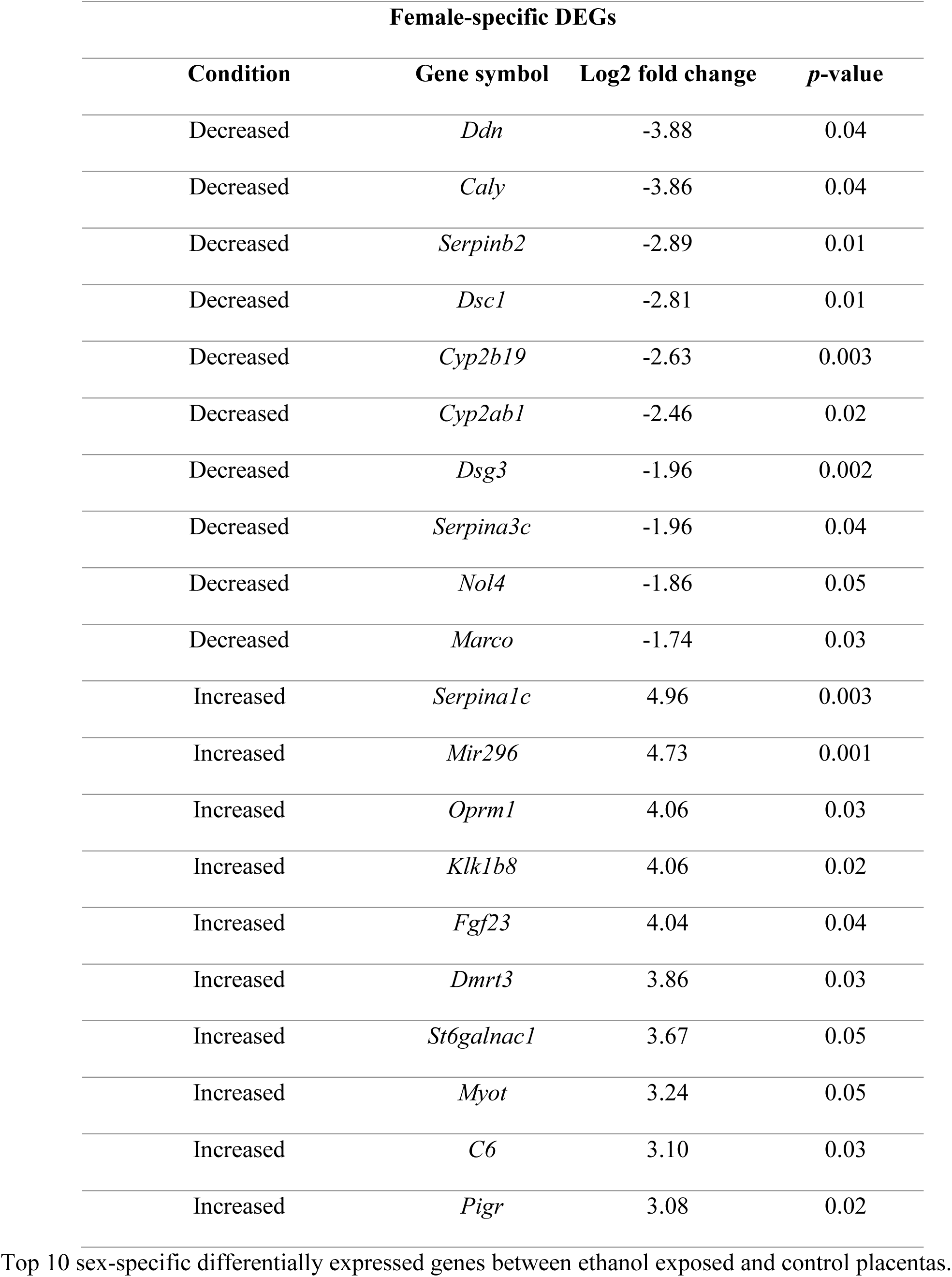
Top 10 decreased and increased sex-specific DEGs.

Functional enrichment analyses of male- and female-specific DEGs revealed divergent pathways. For instance, male DEGs were related to cell migration (*Aqp1*, *Rhoc*), cytoskeletal organization (*Apoa1*, *Bmp10*), cell-substrate adhesion (*Adam8*, *Utrn*), cell junction organization (*Ank6*, *Cdh6*), and blood vessel development (*Flt4*, *Slc16a1*) (Fig. 4E); whereas the female DEGs showed enrichment in pathways involved in cell adhesion (*Itgb3*, *Ret*), serotonin uptake (*Slc22a2*, *Snca*), PIWI-interacting RNA metabolism (*Fkbp6*, *Piwil2*), megakaryocyte development and platelet production (*Gata2*, *Gata4*), and locomotory behavior (*Chrna4*, *Efnb3*) (Fig. 4F). Examples of male- and female-specific DEGs are shown in Figure 4G and 4H.

Notably, we identified 4 genes (*Bahcc1*, *Bhlhe41*, *Prkd3*, *Slc4a1*) with differential expression in the male EtOH-placenta and 2 genes (*Bhlhe41, Slc9a8*) with differential expression in female EtOH-placenta. These genes were among the 40 that were also dysregulated in the placentas of women who reported heavy drinking during pregnancy (Deyssenroth et al., 2024). Amid these, *Bahcc1*, involved in chromatin binding, is linked to KBG syndrome, which is characterized by distinctive craniofacial features, developmental delay, and intellectual disability. Additionally, *Bhlhe41*, which plays a role in regulating circadian rhythms and cell differentiation, was altered in both sexes.

### 3.6. Shared gene expression changes in late-gestation placentas after early preimplantation alcohol exposure

We also identified 54 genes (Table 4) that showed similar differential expression in both male and female ethanol-exposed placentas compared to control following early preimplantation alcohol exposure. For 93% of genes (50/54), we observed comparable changes between the sexes (Fig. S6A-C); however, 4 genes showed divergent changes in males and females, including *Ifit2* (response to interferon-alpha) and *Krt14* (cytoskeleton organization) (Fig. S6D). For *Ifit2*, the control expression level also differed between males and females (Fig. S6D).

**Table 4:**
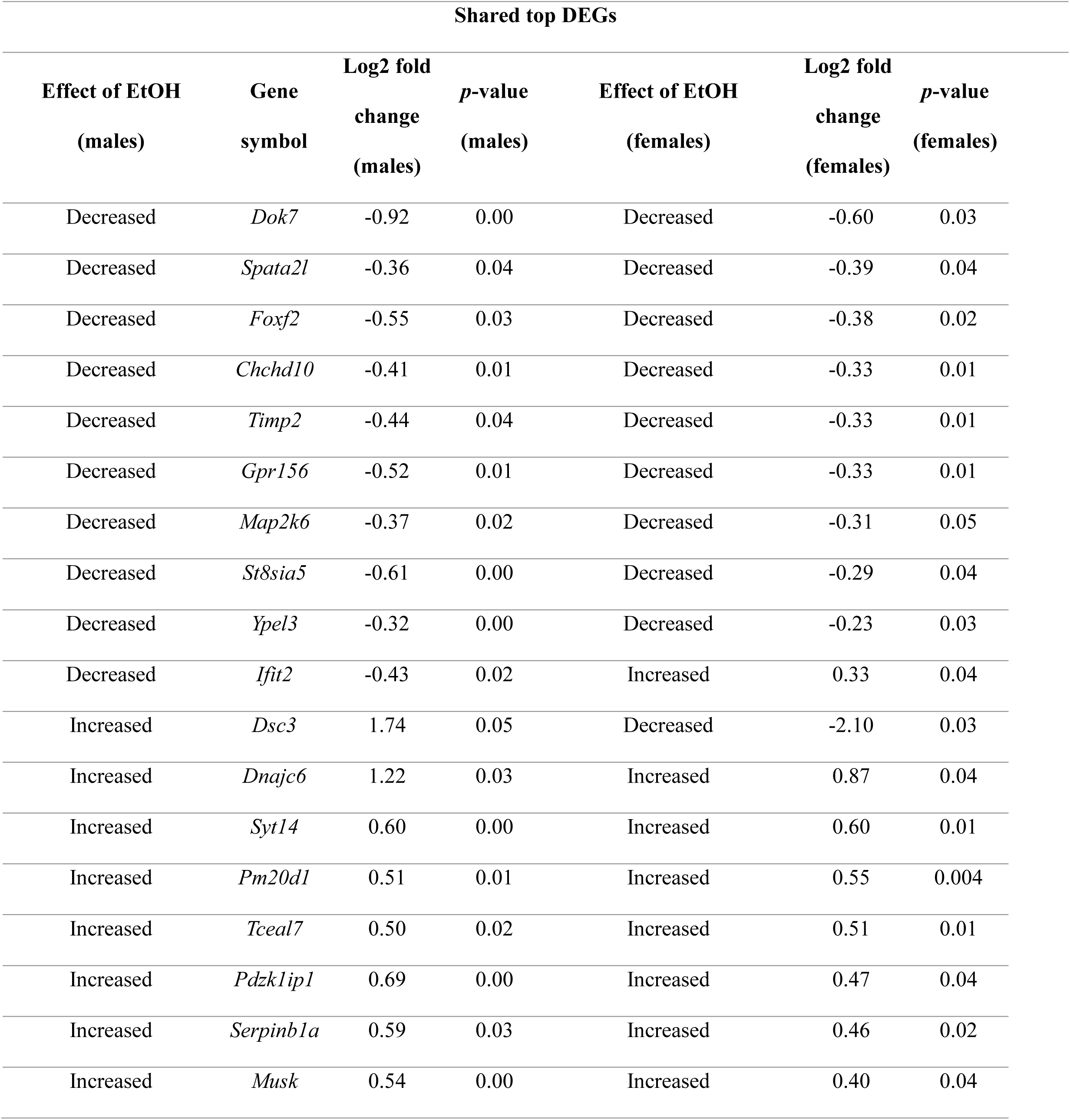
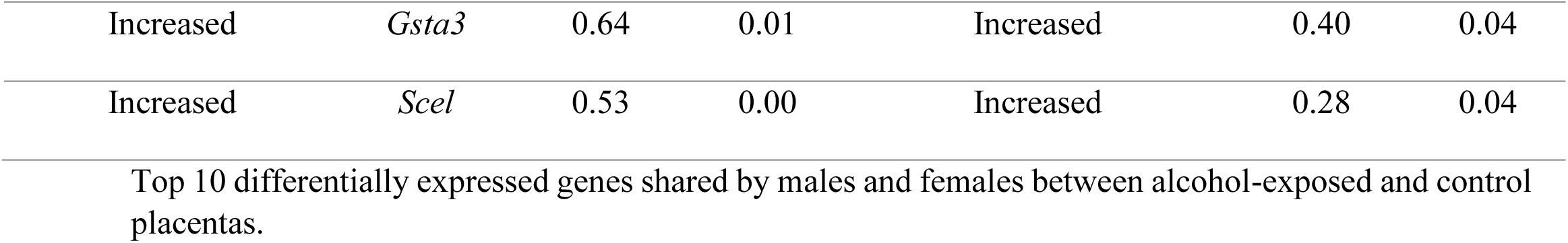
Top 10 shared DEGs between male and female placentas.

Pathway enrichment analysis of these shared DEGs revealed enrichment in neuromuscular junction development (*Musk*, *Pmp22*), sphingolipid biosynthesis (*B3gnt5*, *St8sia5*), artery development (*Chd7*, *Luzp1*), and matrix organization (*Foxf2*, *Kif9*) (Fig. S6E). Additional interesting shared DEGs included *Bhlhe41* (associated with muscle development) and *Cited2* (involved in decidualization, placenta, cardiac, and cortex development), which both showed decreased expression in ethanol-exposed placentas, and *Scel* (which regulates Wnt signaling) and *Syt14* (involved in synaptic transmission), which exhibited increased expression (Fig. S6F). These specific DEGs highlight potential molecular mechanisms and pathways affected by preimplantation alcohol exposure in the placenta regardless of sex.

These findings provide insights into the biological processes that may be impacted by preimplantation alcohol exposure in both male and female placentas, suggesting potential negative consequences of placental dysfunction on embryonic development following early PAE (Legault et al., 2021).

### 3.7. Early preimplantation alcohol exposure does not strictly impact genes crucial for placental development or placental imprinting genes at late gestation

To evaluate the impact of early preimplantation alcohol exposure on genes crucial for placental development, we analyzed our DNA methylation and gene expression datasets using previously published curated gene lists (Legault et al., 2024).

Of the 205 curated placental genes, 193 were present in our DNA methylation dataset, accounting for 7327 tiles located in promoters and intragenic regions (Fig. S7A). However, only six contained DMRs: three in males (*Epas1*, *Id1*, and *Peg10*) and three in females (*Flt1*, *Ppard*, and *Serpine1*). All 205 curated placental genes were included in our mRNA-Seq dataset, of which 19 and 8 were dysregulated following ethanol exposure in males and females, respectively (Fig. S7B). Only one DEG, *Cited2*, was changed in both male and female placentas (Fig. S6B, S6C, and S6F and Fig. S7B). Additionally, *Peg10*’s intronic DMR, which overlaps with the *Peg10* imprinting control region (ICR), exhibited increased DNA methylation and increased expression in male ethanol-exposed placentas (Fig. S7A-B).

To further investigate the impact of preimplantation alcohol exposure on imprinted genes, we searched our datasets for a curated list of 105 genes known to be regulated by genomic imprinting in the placenta (Coan et al., 2005; Golding et al., 2011; Lefebvre, 2012; Okae et al., 2012; Proudhon and Bourc’his, 2010; Wang et al., 2011). In male ethanol-exposed placentas, we identified 17 DMRs located in the intragenic regions of 11 genes (Fig. S8A, S8C). Of these, 12 DMRs in 7 genes were located directly in the ICR. Furthermore, we observed 18 imprinted DEGs in male placentas (Fig. S8B). In contrast, female ethanol-exposed placentas showed fewer DNA methylation changes on imprinting genes, with only 9 DMRs in 9 different genes. Of these, only *Slc22a2* showed differential expression (Fig. S8A, S8B, Table S2 and S3). Detailed examples of DMRs (*Airn*, *Commd1*, *Grb10* and *Peg10*) and DEGs (*Grb10*, *Igf2r*, *Peg10*, *Zrsr1* and *Slc22a2*) for male and female placentas are shown in Figure S8C and S8D.

These results reveal that although several placental and imprinted genes exhibited dysregulated DNA methylation levels or expression following early preimplantation alcohol exposure, the dysregulation was not widespread. These findings align with our morphometric analyses, which did not reveal any significant placental malformations.

### 3.8. Sex-specific regions with both DNA methylation and gene expression changes in the late-gestation following early preimplantation alcohol exposure

To explore the relationship between dysregulated DNA methylation and gene expression profiles in the placenta after preimplantation alcohol exposure, we next compared our lists of promoter and intragenic DMRs and DEGs by examining the expression profiles of the genes associated with 607 male DMRs (542 unique genes), 821 female DMRs (749 unique genes), and 12 shared DMRs (11 unique genes).

Of the 11 genes with shared DMRs, only *Clstn3* and *Tro* exhibited gene expression changes in male ethanol-exposed placentas (Fig. S9A-B), and no expression changes were observed in females. In males, 40 unique genes containing 46 DMRs displayed both DNA methylation and gene expression changes in ethanol-exposed placentas (Fig. 5A). The majority of these DMRs (36; 78%) were associated to small but significant gene expression changes (Fig. S10A). Notably, altered regions/genes included *Apoc1* (lipidic activity), *Carmil1* (implicated in actin filament processes), *Slc16a3* (encodes the MCT4 protein, important for transmembrane transport, placental organization, and vascularization) and *Zfhx3* (a transcription factor important for development) (Fig. 5B).

**Figure 5:**
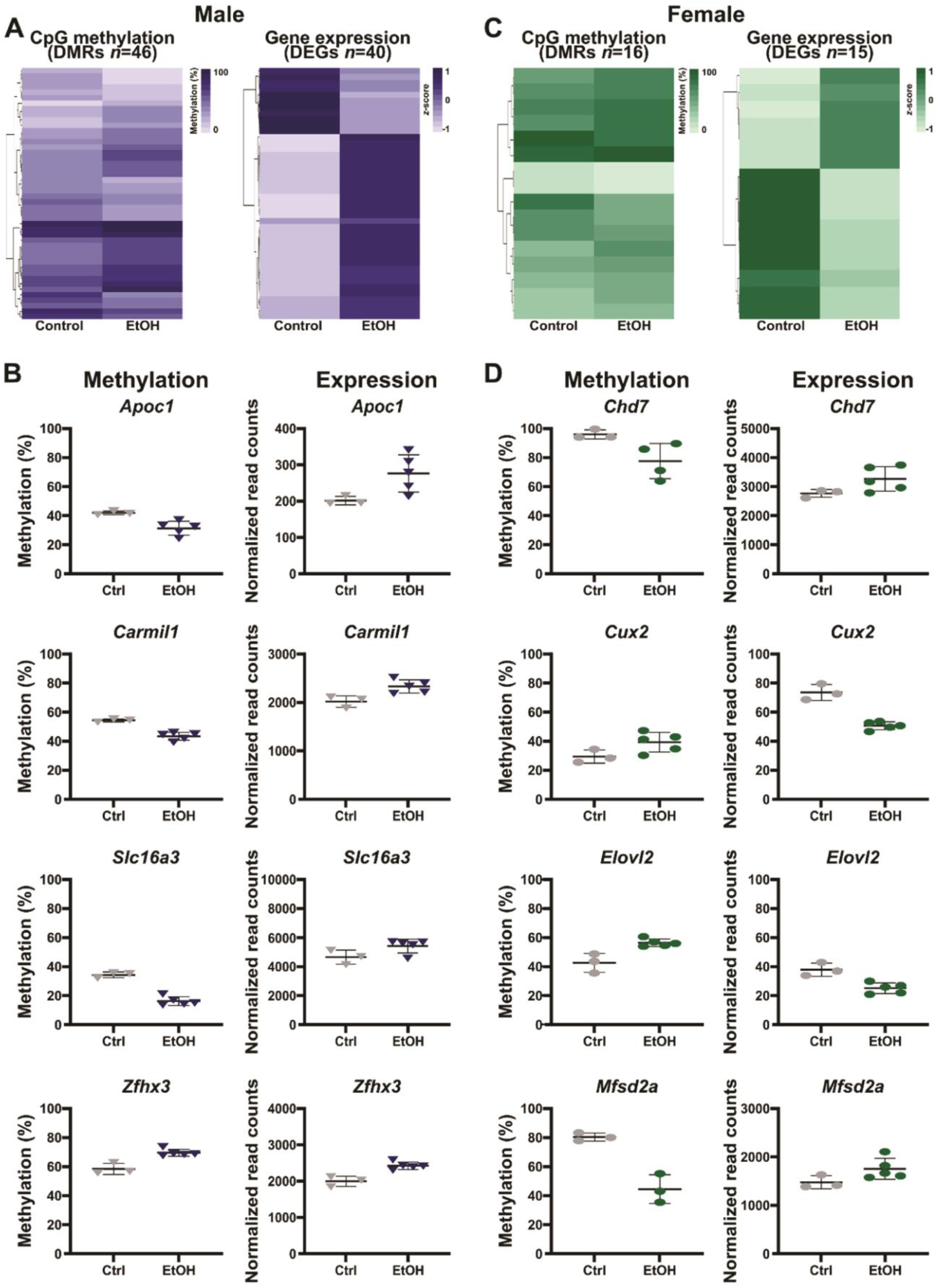
Sex-specific regions show both DNA methylation and gene expression alterations in the placenta at late gestation in response to early preimplantation alcohol exposure. **A)** Heatmaps of CpG methylation (left) and gene expression (*z*-score; right) levels of DEGs containing intragenic DMRs between control and ethanol-exposed male (46 DMRs, 42 DEGs) **B)** Normalized read counts of DEGs and DMRs in male placentas. **C**) Heatmaps of CpG methylation (left) and gene expression (*z*-score; right) levels of DEGs containing intragenic DMRs between control and ethanol-exposed female (16 DMRs, 15 DEGs) placentas. **D**) Normalized read counts of DEGs and DMRs in female placentas. Data represent the mean ± SD.

In females, 15 unique genes containing 16 DMRs showed both differential methylation and gene expression changes in ethanol-exposed placentas (Fig. 5C). Eleven of these DMRs (69%), associated to 7 genes (47%), revealed changed expression by (Fig. S10B). Noteworthy genes included *Chd7* (implicated in cardiac, muscle, and neuronal development), *Cux2* (related to neuronal pathways and functions, including cortical interneuron development), *Elovl2* (involved in fatty acid processes), and *Mfsd2a* (implicated in the establishment of the blood-brain barrier) (Fig. 5D).

These results offer significant insights into the enduring effects of preimplantation alcohol exposure on multiple genes, shedding light on the potential interplay between gene expression and DNA methylation. The findings underscore the importance of understanding sex-specific DNA methylation dysregulation as a potential mechanism shaping the distinctive gene expression patterns observed in both male and female placentas.

### 3.9. Bulk RNA-seq deconvolution reveals no distribution shift in placental cell subtypes following preimplantation alcohol exposure

Although our morphological analyses indicated that early alcohol exposure did not alter the overall morphometric structure of the placenta (Fig. S1), we aimed to confirm that the observed molecular changes (DNA methylation and gene expression) were directly affecting the cells themselves, rather than being influenced by alterations in placental cell composition. To do so, we used single-cell RNA-seq data from E18.5 mouse placenta (Fu et al., 2024) to determine cell type proportions through a deconvolution approach (Fig. S11). The estimated cell type proportions remained constant across all samples, with no significant differences observed in both male and female samples (Table S4).

### 3.10. Unveiling predictive molecular signatures in the placenta for early preimplantation alcohol exposure

Next, we wanted to determine which molecular signature, either DNA methylation or gene expression profiles, proves superior in predicting early preimplantation alcohol exposure in the mouse placenta. We used a random forest classifier and least absolute shrinkage and selection operator (LASSO) regression, a machine-learning technique, to create a predictive model using identifiable markers found in our transcriptomic or DNA methylation data. To enhance accuracy and effectiveness, we trained the model using both male and female samples together, increasing our statistical strength and refining the analytical process.

First, we used gene expression profiles and performed LASSO regression to establish specific transcriptomic signatures for ethanol-exposed placentas. Despite being statistically significant, the 20 DEGs identified were not different enough between the conditions to establish a good transcriptomic signature of preimplantation alcohol exposure in late-gestation placentas, with small Euclidean distances between samples of the two conditions and poor statistical coefficient of importance (Fig. S12A-C).

Interestingly, LASSO regression using our DNA methylation profiles identified a set of 24 statistically significant DMRs that formed a specific signature enabling the effective discrimination of ethanol-exposed samples from controls for both principal component analysis and Euclidean distance calculations (Fig. 6A-B). These DMRs were both intragenic (Breton-Larrivée et al., 2023) and intergenic (Cole, 2009), and each had different levels of importance based on their statistical parameters, notably the LASSO coefficient, Wilcoxon *p*-value, and AUC generated by LASSO analysis (Fig. 6C). The most significant DMR in the signature was located in an intergenic region close to *Nup35*. Additionally, there were DMRs in genes such as *B3galt1*, *Kcng1*, *Mir3960*, *Mir6940*, *Psd3*, and *Sesn1*, along with intergenic DMRs close to *Bicd1*, *Gatsl2*, and *Zeb2* (Fig. 6C-D).

**Figure 6:**
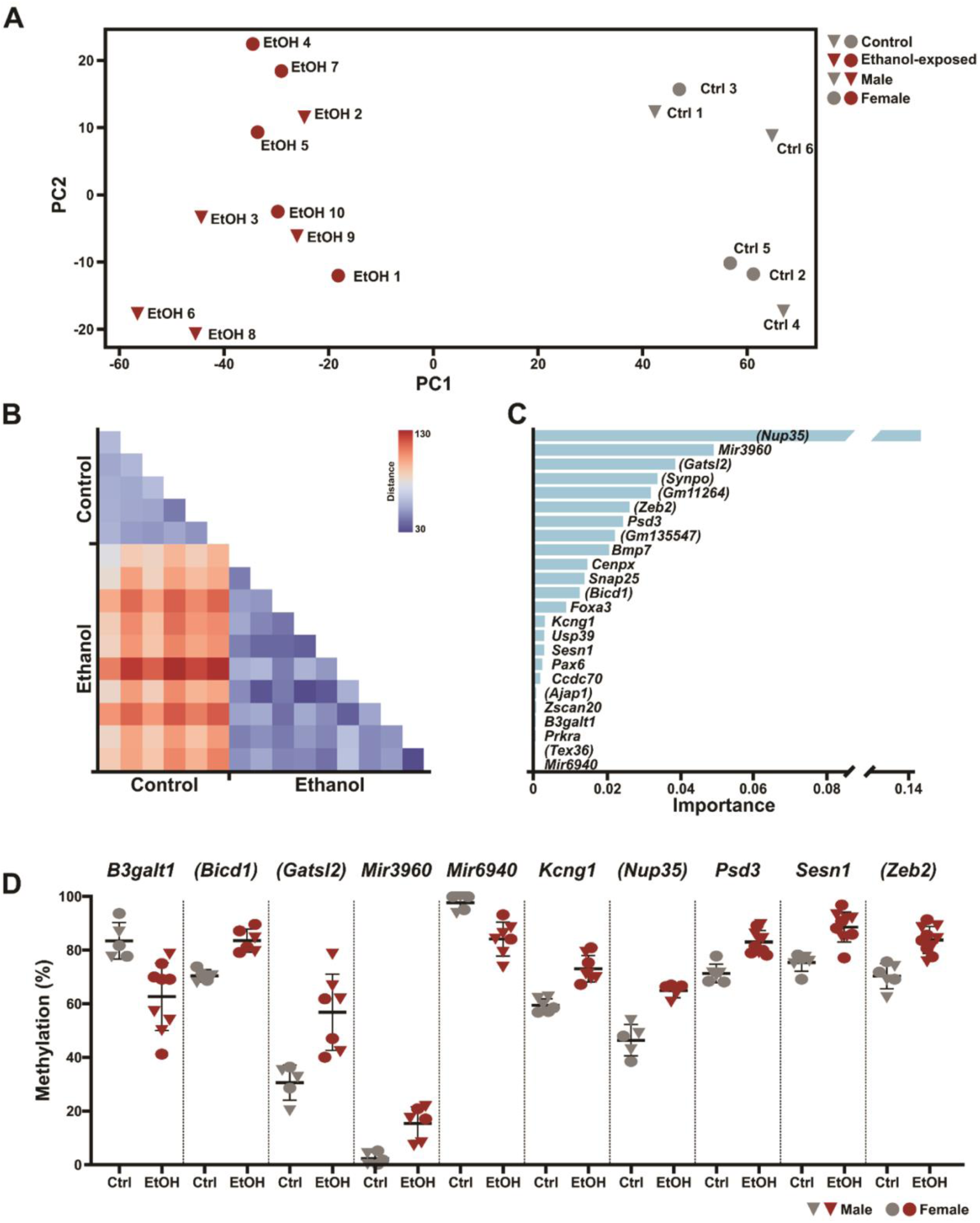
Identification of distinctive placental DNA methylation signatures indicative of early preimplantation alcohol exposure. **A)** Principal component analysis showing the clustering of control and ethanol-exposed samples based on their CpG methylation levels in the 24 DMRs making up the signature.. **B)** Heatmap of the Euclidean distances of each control and ethanol-exposed sample based on their CpG methylation levels in the 24 DMRs. **C)** Feature importance plot of the DMRs based on LASSO analysis. **D)** CpG methylation levels in control and ethanol-exposed male and female placental DMRs making up the signature. Genes in brackets are the closest gene to the intergenic DMR. Ctrl: control, EtOH: ethanol-exposed.

These findings unveil a distinctive molecular signature in placental tissue, highlighting a reliable DNA methylation pattern that serves as a robust molecular tool for detecting early embryonic alcohol exposure at birth.

## 4. DISCUSSION

In this study, we examined the molecular and developmental effects of PAE during early preimplantation on the late-gestation placenta. Our analyses are the first to reveal extensive sex-specific alterations in both DNA methylation and gene expression patterns in late-gestation mouse placentas exposed to ethanol. Interestingly, despite these molecular changes, the overall morphological development of the placenta appeared unaffected. Finally, employing machine learning, we demonstrate that DNA methylation profiles in the placenta emerge as more robust indicators of early alcohol exposure compared to gene expression signatures.

### 4.1. Sex-specific impacts of preimplantation alcohol exposure on embryo growth and related pathways

Prenatal alcohol exposure was previously shown to alter embryonic development in different contexts, by inducing fetal growth restriction or growth delay (Breton-Larrivée et al., 2023; Legault et al., 2021). With our model, we found that the impact on growth and related pathways was more pronounced in male ethanol-exposed placentas, consistent with the observed male-specific changes in genes regulating growth, cytoskeletal organization, placental development, and vascularization. Furthermore, we noted a reduction in embryo weight only in male, indicating that males may be more susceptible to growth delays following preimplantation alcohol exposure. However, it is worth noting that in our previous studies using this model, we observed similar percentages of growth-delayed embryos (∼5%) between males and females at mid-gestation (Legault et al., 2021). It is possible that some female embryos with growth delays at mid-gestation may catch up later in development, eventually achieving a similar embryo weight to control. Several studies have demonstrated that female embryos generally exhibit greater adaptability to adverse *in utero* environments than male embryos, facilitating their development even in the presence of altered resource availability (Bale, 2016; Christians et al., 2023; Meakin et al., 2021). Male ethanol-exposed placentas displayed altered methylation and/or expression patterns in several genes that are directly linked to embryonic growth restriction. For example, a study recently showed that injecting *fibroblast growth factor* 2 (*Fgf2;* downregulated in male ethanol-exposed placentas) into a rat model of preeclampsia improved clinical outcomes, including attenuating fetal growth restriction (Martinez-Fierro et al., 2022). Mutation of *Fgf1* (increased methylation in male ethanol-exposed placentas) has been linked to preeclampsia in humans (Marwa et al., 2016), while *Bmp6* (decreased methylation in male ethanol-exposed placentas) was downregulated in the fetal kidneys of rats with intrauterine growth restriction in a model of induced hypertension due to chronic insulin administration (Bursztyn et al., 2006). Changes in these genes may significantly contribute to the inability of male embryos to develop normally and achieve the embryo weights of control embryos.

### 4.2. Distinct patterns of DNA methylation dysregulation in female and male ethanol-exposed placentas

DNA methylation analysis of late-gestation placentas following preimplantation alcohol exposure uncovered sex-specific DMRs. Moreover, we also observed sex-specific alteration patterns in DNA methylation changes as well as in the proximity of the sex-specific DMRs to CpG-rich regions, suggesting a sex-specific effect on DNA methylation based on CpG content following preimplantation alcohol exposure.

In a recent study, we discovered genome regions with sex-biased DNA methylation profiles in control placentas at E18.5 (Legault et al., 2024). When we compared the DMRs observed following preimplantation alcohol exposure with these sex-biased regions, we found that 115 male DMRs and 188 female DMRs were also sex-biased at late gestation. Importantly, these ethanol-exposed DMRs and sex-biased regions exhibited distinct CpG distributions based on sex. Notably, we observed a higher proportion of female-specific DMRs located in sex-biased regions in CpG islands and shores than male-specific DMRs located in sex-biased regions (Legault et al., 2024). These differentially methylated regions between male and female embryos could help explain the sex-specific changes observed in both DNA methylation and gene expression. It is known that sex-biased differences exist during early development, and that divergent DNA methylation profiles can contribute to sex-specific DMRs in late-gestation placentas. One hypothesis is that these specific regions are more vulnerable to harmful exposures from the maternal environment due to their sex-specificity during development. Epigenetic dysregulation in these specific regions may progress differently in males and females, resulting in sex-specific DMRs in the late-gestation following preimplantation alcohol exposure.

Our findings highlight the value of exploring placental molecular pathways in the context of FASD. DMRs and DEGs in our model showed alterations with genes recently identified as deregulated in the placentas of women who engaged in heavy drinking during pregnancy, including the crucial 3-week period around conception (Deyssenroth et al., 2024). The variations in alcohol exposure paradigms (single vs. multiple exposures), the influence of additional factors such as smoking (Hoch et al., 2023), sex-specific analyses, differences in exposure timing (pre-implantation vs. variable developmental time points), and species-specific differences may account for the limited overlap between our study and that of Deyssenroth *et al*. Nonetheless, these observations underscore the relevance of our approach and its potential for furthering understanding in this field. Further studies investigating dysregulated DNA methylation in the trophectoderm and placenta during early development could provide insights into the variability of these sex-specific regions after preimplantation PAE. Understanding the dynamics and mechanisms underlying these sex-specific effects will contribute to our knowledge of how PAE impacts placental function and subsequent developmental outcomes in a sex-dependent manner.

### 4.3. Preimplantation alcohol exposure had a greater impact on imprinted and placental development genes in male placentas

The methylation patterns of the ICRs of imprinted genes and the DNA methylation and expression patterns of genes crucial for placental development can significantly influence proper embryo development (Hemberger and Cross, 2001; Plasschaert and Bartolomei, 2014). While placental development being largely completed by late gestation, epigenetic and transcriptomic dysregulation of key genes can disrupt its function, especially when the dysregulation occurs at earlier developmental stages (Apicella et al., 2019). Although histological analysis of the junctional and labyrinth zones showed comparable results between ethanol-exposed and control placentas in both sexes, we observed more changes in imprinted and placental development genes in male ethanol-exposed placentas compared to females.

Among the dysregulated genes, we observed differential methylation and expression of the imprinted gene *Grb10* in male ethanol-exposed placentas. *Grb10* encodes a growth restrictor, and irregular imprinting, especially in its ICR, can directly affect embryonic development (Plasschaert and Bartolomei, 2014). The increased *Grb10* expression observed in male ethanol-exposed placentas could therefore directly impact embryonic growth and development. Additionally, we observed dysregulation in both DNA methylation and gene expression in the *H19*/*Igf2* cluster, including increased expression of *Igf2r* in male ethanol-exposed placentas and increased methylation in the ICR of *Airn* (antisense of *Igf2r*).

Like *Grb10*, *Igf2r* negatively regulates embryonic growth and development (Apicella et al., 2019; Hemberger and Cross, 2001; Plasschaert and Bartolomei, 2014; Sleutels et al., 2003). This imprinted cluster has been previously identified using a similar paradigm of alcohol exposure during preimplantation. In their study, Haycock and Ramsay showed decreased DNA methylation in the ICR of *H19* in the mid-gestation (Haycock and Ramsay, 2009). They also observed significant decreases in the weights of both ethanol-exposed embryos and placentas at mid-gestation and found a strong correlation between DNA methylation of the paternal *H19* allele and weight loss in the ethanol-exposed group (Haycock and Ramsay, 2009). The abnormal methylation and expression profiles of imprinted genes collectively impact proper embryonic development, even though placental morphology is not affected.

Analysis of placental development genes also revealed more genes with altered expression profiles in male ethanol-exposed placentas. Notably, we observed increased expression of *Dnmt3l*, a coactivator of *Dnmt3a* and *Dnmt3b*, which catalyze *de novo* DNA methylation (Bourc’his et al., 2001). During placental development, *Dnmt3l* is essential for establishing proper maternal gene imprinting profiles and is highly expressed in trophoblasts (Bourc’his et al., 2001; Joh, 2018; Logan et al., 2013). While the impact of *Dnmt3l* overexpression on the placenta is unknown, studies have demonstrated morphological defects and abnormal trophoblast patterns in *Dnmt3l* knockout mice, providing further evidence of its critical role in proper embryonic development (Andrews et al., 2023; Arima et al., 2006). We cannot exclude the potential impact of dysregulated *Dnmt3l* expression on the irregular DNA methylation and transcriptomic profiles observed in certain imprinted genes, such as the maternally methylated genes *Peg10*, *Jade1*, or *Zrsr1,* in male ethanol-exposed placentas.

### 4.4. Dysregulation of the placental epigenetic and transcriptomic landscape could play a role in proper embryonic brain development

The placenta-brain axis has recently received considerable attention, and it is now well-established that placental dysregulation can lead to adverse effects on brain development and function (Bale, 2016; Rosenfeld, 2021). Neurodevelopmental disorders and intellectual disabilities are commonly observed in FASD, resulting from disruptions during brain development and incomplete maturation (Popova et al., 2023). While the impact of endocrine placental dysfunction on brain development is increasingly understood, the role of placental epigenetic and transcriptomic dysregulation has been less explored. Among the many enriched pathways observed in our datasets, two main pathways emerged that could directly affect normal embryonic brain development.

First, we found that male-specific DMRs were enriched for components of a pathway related to synaptic transmission, with dysregulated genes including *Asic1*, *Cbln1*, and *Plk2*. *Asic1* and *Cbln1* are required for normal neuronal development (Lim et al., 2021; McCormick and Gupton, 2022), and *Plk2* has been linked to epilepsy because of its role in synaptic potentiation (Seeburg and Sheng, 2008). Additionally, our transcriptome analysis revealed that one of the main pathways enriched in female ethanol-exposed placentas was related to serotonin metabolism. Serotonin is a neurotransmitter that regulates embryonic neurodevelopment, and the placenta plays a neuroendocrine role by producing and providing serotonin to the developing forebrain (Rosenfeld, 2020). Placental insufficiency, including failing to provide an adequate serotonin supply to the embryo, is associated with the development of mental health issues and reduced cognitive abilities (Rosenfeld, 2020). Knockout of *Itgb3*, which was downregulated in female ethanol-exposed placentas, had adverse effects on the placenta’s vascularization structure in a sheep model, compromising nutrient transport (Frank et al., 2021). ITGB3 is also directly involved in serotonin uptake by interacting with different serotonin transporters (Whyte et al., 2014). Interestingly, *Slc22a2*, a gene implicated in serotonin clearance has been identified as overexpressed in female ethanol-exposed placentas (Bacq et al., 2012). Together, changes in these two genes could negatively modulate serotonin uptake in the developing brain *via* altered placental endocrine function and disrupted communication, further emphasizing the impact of the placenta on embryonic health and development.

### 4.5. Dysregulated placental DNA methylation profiles serve as a robust signature of early preimplantation alcohol exposure

Our study unequivocally demonstrates that alcohol exposure as early as the 8-cell stage induces enduring molecular alterations in both DNA methylation and gene expression profiles in the mouse placenta. However, we show that the stability and persistence of DNA methylation alterations across ethanol-exposed placenta profiles underscore its potential superiority as a molecular signature for capturing and reflecting the long-term impact of alcohol exposures on developmental processes. This further supports the ability to employ DNA methylation patterns as a viable, reliable, and precise indicator of previous alcohol prenatal exposure.

In recent years, two research groups showed DNA methylation changes in buccal epithelial cell swabs from children with confirmed FASD (Laufer et al., 2015; Lussier et al., 2018; Portales-Casamar et al., 2016). Moreover, one of these studies applied a machine learning approach with the identified epigenetic signature to train and further efficiently validate the identification of confirmed FASD cases (Lussier et al., 2018). Although these cells are easily accessible and require no invasive procedure to obtain, their epigenome will vary with age and may generate some bias in the generation of accurate DNA methylation signatures. Moreover, the high renewal rate of these surface cells may not homogeneously reveal PAE-induced epigenetic modifications. PAE’s heterogenous paradigms and their effects on the epigenomes of buccal epithelial cells are also an obstacle in establishing a universal DNA methylation signature. Tissues or cell types more directly exposed to alcohol during gestation may be more promising alternatives. In this perspective, the placenta could be an effective means to detect PAE at birth since its development occurs rapidly during gestation. A previous study using preconception and gestational cannabis also observed shared changes in DNA methylation in the placentas and brains of exposed embryos (Laufer et al., 2022), indicating that the placenta may be able to effectively mirror some DNA methylation alterations present in the brain. The short lifespan of the placenta (resulting in minimal cell renewal) makes it an optimal candidate tissue to profile epigenetic alterations caused by PAE any time during gestation. The placenta can be thought of as an organ having a high potential to teach us more about what transpired during pregnancy and how it may have affected embryonic development.

As mentioned earlier, the absence of a molecular diagnostic tool for PAE and FASD results in most cases going undetected until children reach school age, where noticeable neurodevelopmental symptoms emerge (Popova et al., 2023). While our study used a limited number of mouse samples, the application of DNA methylation profiles and machine-learning approaches has enabled us to identify robust molecular signatures for early embryonic alcohol exposure. Even though this molecular signature has yet to be validated, it serves as a proof of concept that could be applied to humans, enabling early diagnosis and optimal support for newborns with FASD from their early days. Future research should investigate multiple time points using whole-genome DNA methylation approaches combined with single-cell sequencing technologies in the same mouse conceptus (placenta and embryonic cells) and human placental tissues. This strategy will yield a deeper understanding of early embryonic alcohol exposure, providing enhanced datasets for machine-learning approaches and ultimately establishing more accurate and comprehensive molecular signatures.

Nevertheless, ethical concerns surrounding PAE diagnostic tests, including the risk of stigmatization and discrimination, challenges in obtaining informed consent, and issues related to accuracy, need careful consideration. Thus, there is a crucial need to balance these concerns with initiatives aimed at enhancing our understanding of the mechanisms and risks associated with alcohol consumption during pregnancy, addressing issues like lack of awareness and unintentional exposure. Additionally, there is a call for the development of clearer guidelines to assist women and healthcare professionals in making informed decisions about alcohol consumption, prioritizing the well-being of both the mother and the child.

## 5. CONCLUSION

In this study, we showed that early preimplantation alcohol exposure can alter the molecular profile of late-gestation mouse placentas. More precisely, our study provides valuable insights into the sex-specific epigenetic and transcriptomic consequences of early PAE on the placenta, contributing to a better understanding of its potential effects on placental function and embryo development. Furthermore, this study highlights for the first time that the placental DNA methylation profiles provide robust molecular signature for early PAE detection. Further studies based on our model will help decipher the mechanisms of sex-specific placental DNA methylation and gene expression changes on embryonic development and their long-term implications.

## DECLARATIONS

### Ethics approval

All animal work conducted in this study was approved by the CHU Ste-Justine Research Center *Comité Institutionnel de Bonnes Pratiques Animales en Recherche* (CIBPAR) under the guidance of the Canadian Council on Animal Care (CCAC).

### Consent for publication

Not applicable

### CRediT authorship contribution statement

**Lisa-Marie Legault:** Writing – review & editing, Writing – original draft, Visualization, Methodology, Investigation, Formal analysis, Conceptualization. **Thomas Dupas:** Writing – review & editing, Writing – original draft. **Mélanie Breton-Larrivée:** Visualization, Methodology, Investigation, Formal analysis. **Fannie Filion-Bienvenue:** Formal analysis. **Anthony Lemieux:** Formal analysis. **Alexandra Langford-Avelar:** Formal analysis. **Serge McGraw:** Writing – review & editing, Supervision, Conceptualization, Funding acquisition. All authors read and approved the final manuscript.

## Data availability

All sequencing datasets generated and used in this study will be publicly available via the Gene Expression Omnibus repository (GSE254847).

## Acknowledgements

We extend our gratitude to Dr. Vincent Couture (Université Laval) for his thoughtful insights in shaping the discussion on ethical concerns, the McGraw lab for their critical comments and suggestions, Dr. Karine Doiron, Elizabeth Maurice-Elder and High-Fidelity Science Communications for editing this manuscript, the staff of the Centre de Recherche du CHU Sainte-Justine animal facility for their assistance, and the Centre d’expertise et de services Génome Québec for their support.

## Appendix A. Supplementary data

The following are the Supplementary data to this article:

Supplementary Data 1: Figures S1-S12, Tables S2 and S3.

Supplementary Data 2: Table S1.

Supplementary Data 3: Table S4.

## SUPPLEMENTAL FIGURES and LEGENDS

**Supplemental Figure 1:**
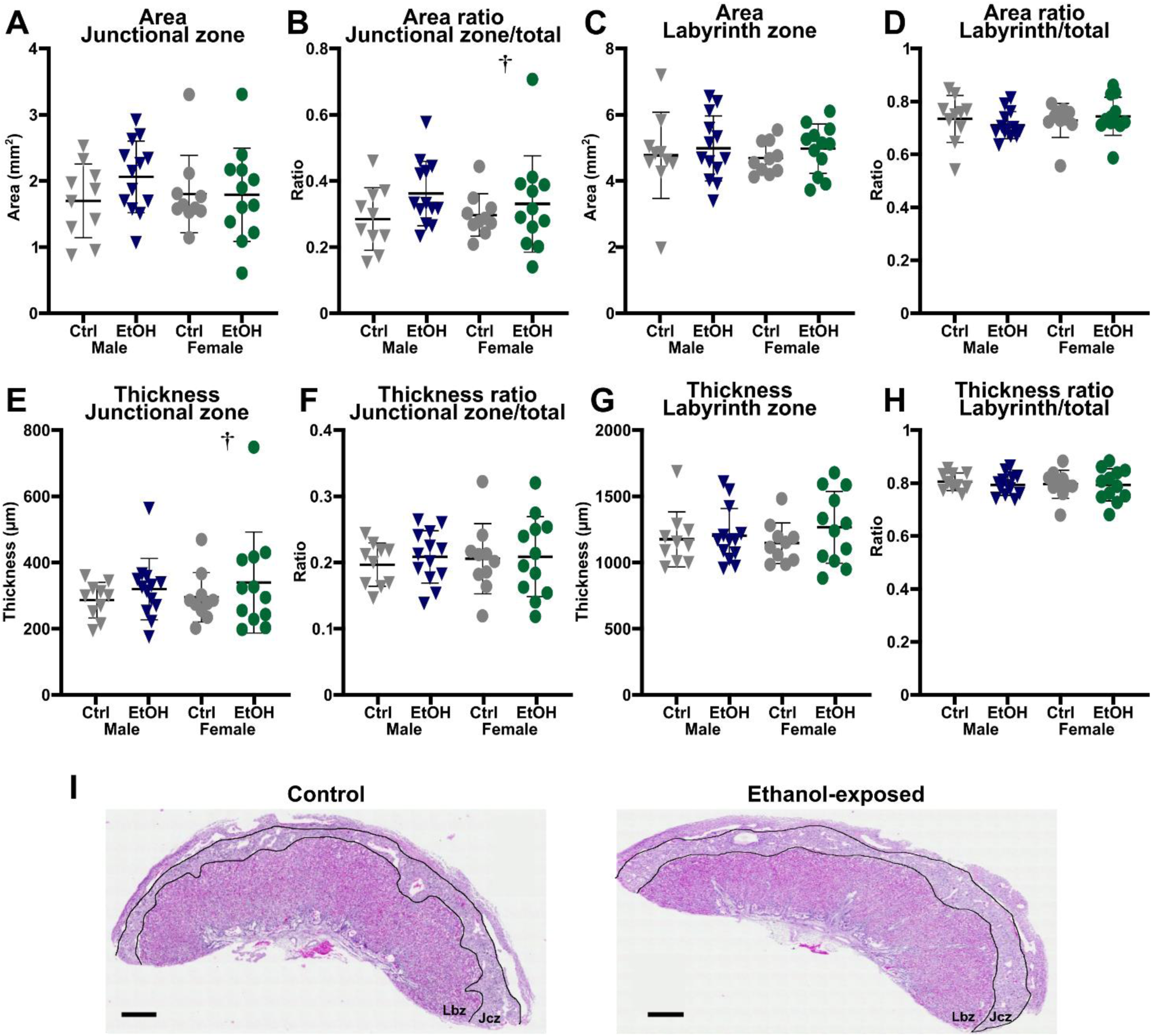
Increased variability in female ethanol-exposed placenta morphometric measurements. Histological morphometric measurements of E18.5 placentas based on placental cross-sections (Males : Ctrl *n* = 10 ; EtOH *n* = 13. Females Ctrl *n*= 10; EtOH *n* =12). Assessment of the impact of preimplantation alcohol exposure on **A)** junctional zone (Jcz) area; **C)** Labyrinth zone (Lbz) area; **E)** thickness of the Jcz; **G)** thickness of the Lbz and their respective ratio to the total placenta area (**B, D**) or thickness (**F, H**). **I)** Representative images of hematoxylin and eosin-stained control and ethanol-exposed samples. The Jcz and Lbz are delimited by black lines. All data represent the mean ± SD. Significant differences were assessed by *t*-test with Welch’s correction and *F* tests for the variance analysis. † Higher variance in ethanol-exposed placentas; *F*-test, *p*˂0.05. Ctrl: control, EtOH: ethanol-exposed.

**Supplemental Figure 2:**
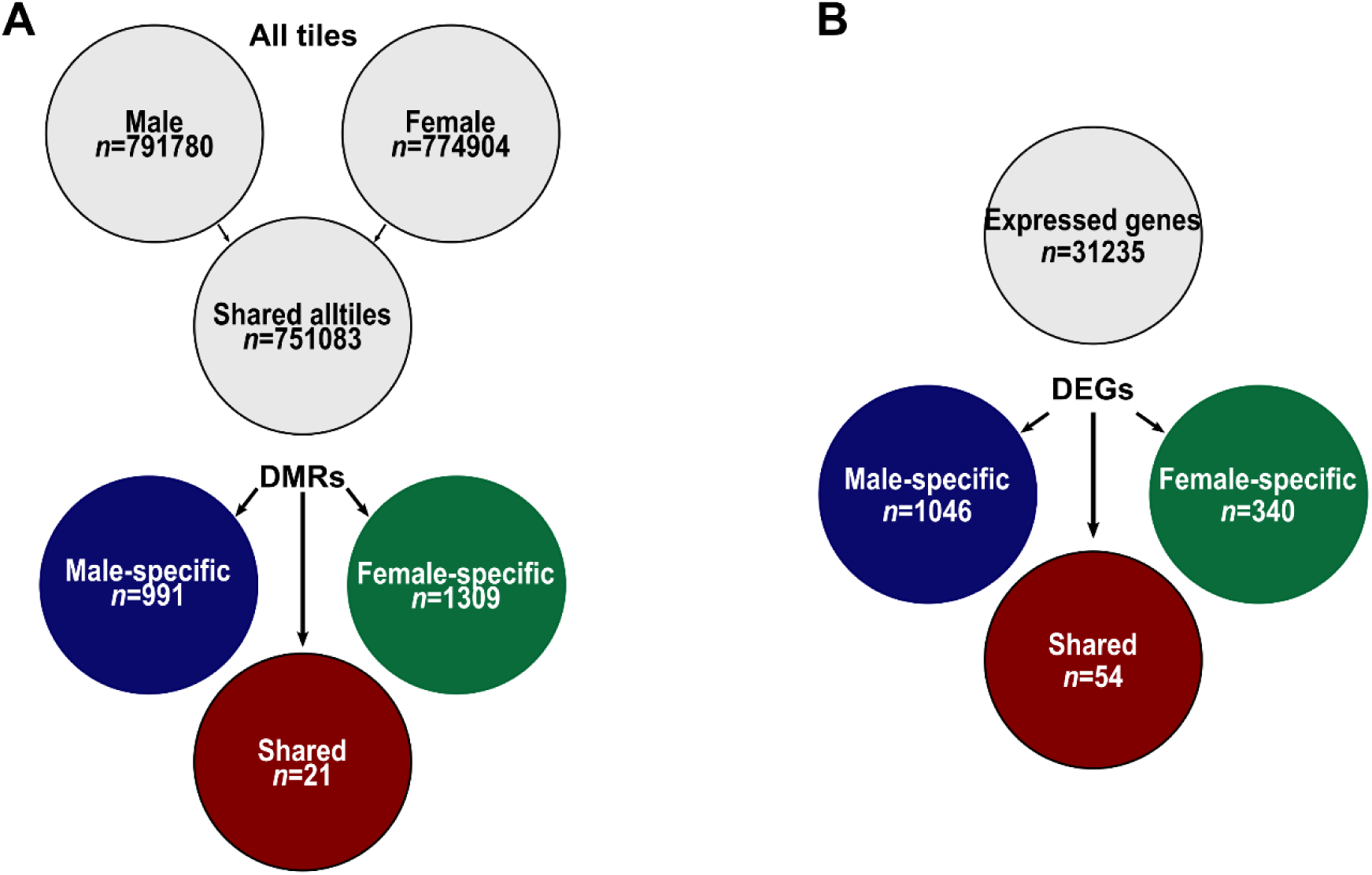
Establishing DNA methylation profiles in alcohol-exposed late-gestation placentas (E18.5) **A)** Schematic of the sex-related genome-wide CpG methylation analysis in male (Ctrl, *n*=3; EtOH, *n*=5) and female (Ctrl, *n*=3; EtOH, *n*=5) placentas at E18.5 to identify all tiles (100bp, ˃ 10× coverage) associated with either males (791780), females (774904), or both sexes (751083), as well as male-specific (991), female-specific (1309), and common (*n*=21) DMRs). **B)** Schematic of sex-related differential transcriptomic analyses in male (Ctrl, *n*=3; EtOH, *n*=5) and female (Ctrl, *n*=3; EtOH, *n*=5) placentas at E18.5. Identification of all expressed genes in both sexes (31,235) and male-specific (1046), female-specific (340), and shared (54) DEGs.

**Supplemental Figure 3:**
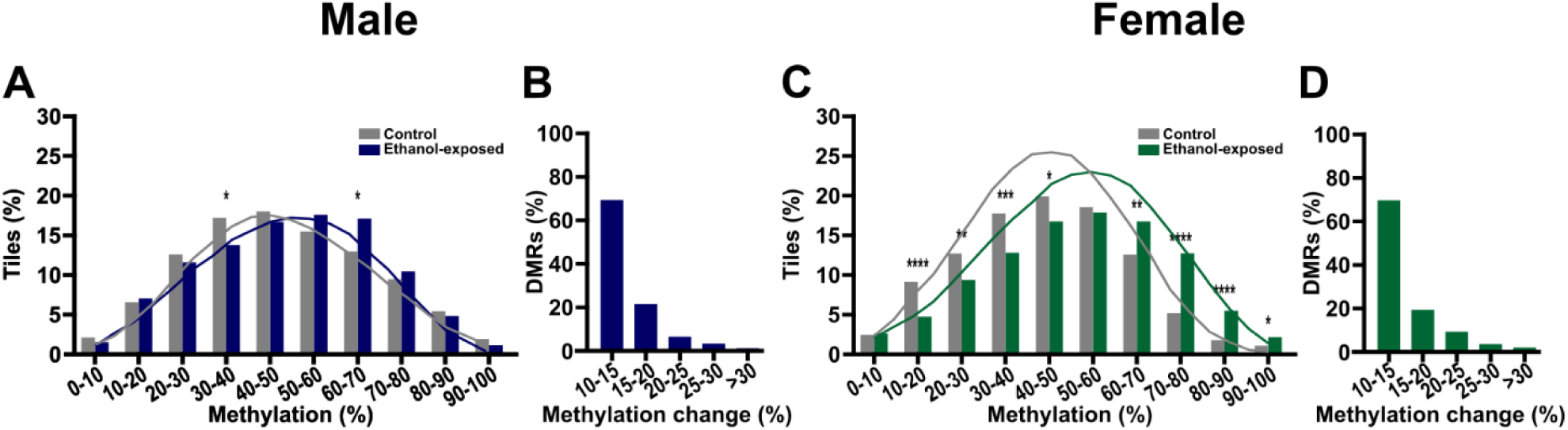
DNA methylation level distribution across differentially methylated regions. Tiles distributions across DNA methylation levels in **A)** male- and **C)** female-specific control and ethanol-exposed DMRs. Proportion of **B)** male and **D)** female DMRs associated with changes in CpG methylation levels between control and ethanol-exposed placentas. Data represent the mean ± SD. *: *p*<0.05, **: *p*<0.01, ***: *p*<0.001, ****: *p*<0.0001 by two-proportions *z*-test.

**Supplemental Figure 4:**
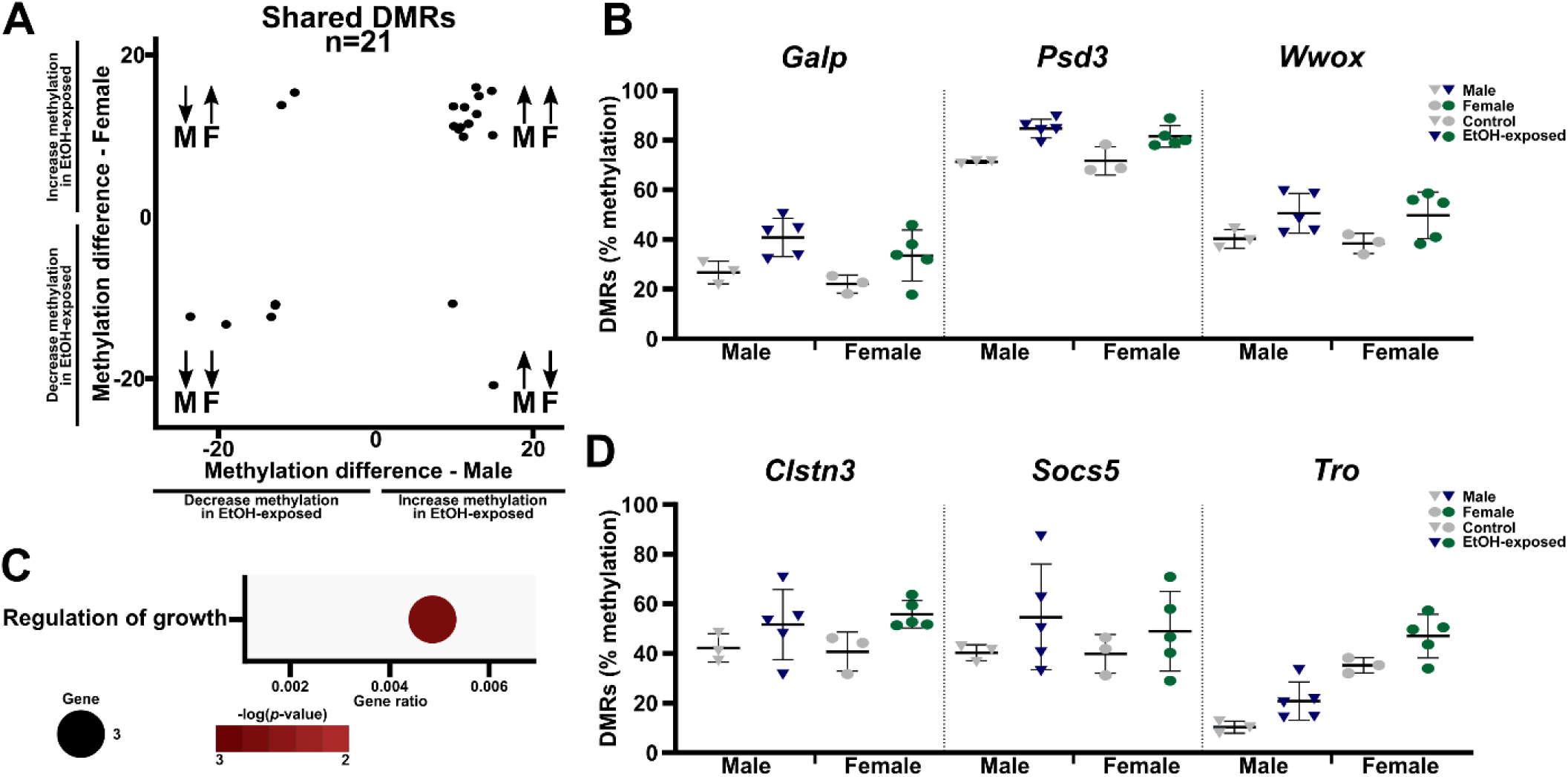
Regions presenting shared differential DNA methylation patterns in ethanol-exposed male and female placentas are enriched in genes related to growth. **A)** Percentage of change in the 21 shared DMRs in males and females. **B)** CpG methylation levels of shared DMRs in control and ethanol-exposed male (left; grey/blue) and female (right; grey/green) placentas. **C)** Top pathways enriched for shared DMRs (11 unique gene DMRs) based on Metascape analysis. **D)** CpG methylation levels for DMRs located in enriched growth-regulating genes (from **C**) in control and ethanol-exposed male (left dots; grey/blue) and female (right dots; grey/green) placentas. Data represent the mean ± SD. Ctrl: control, EtOH: ethanol-exposed.

**Supplemental Figure 5:**
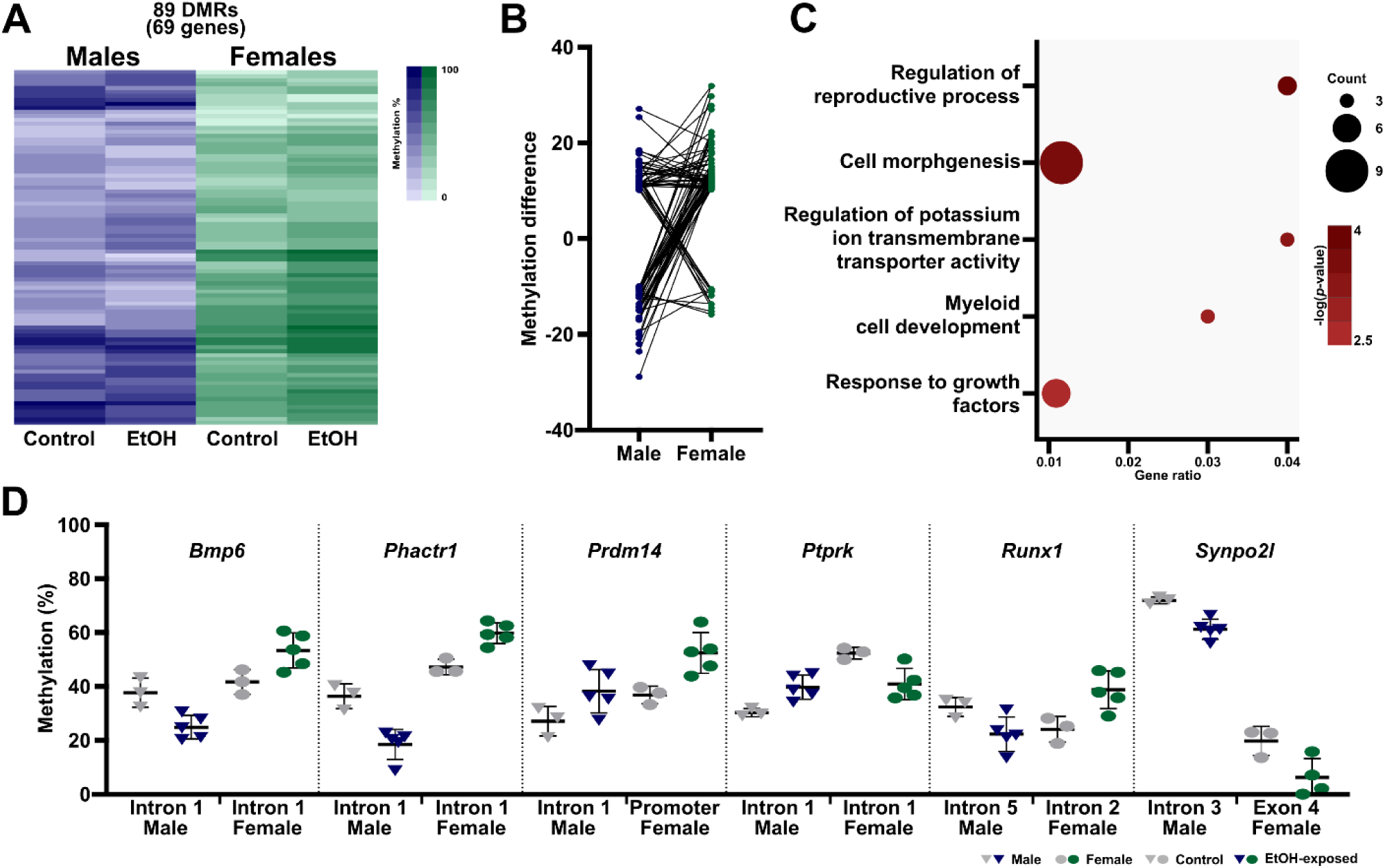
Sex-specific changes in different regions of the same genes in ethanol-exposed placentas. **A)** Heatmaps of the CpG methylation levels of DMRs within the same genes in male (blue) and female (green) placentas. **B)** Methylation differences between males and females. Connected dots represent DMRs in a shared gene. **C)** Top pathways enriched for shared genes with male and female intragenic DMRs (69 unique genes) based on Metascape analysis. **D)** CpG methylation levels of DMRs located in genes related to top enriched pathways in control and ethanol-exposed male (left dots; grey/blue) and female (right dots; grey/green) placentas. Data represent the mean ± SD. Ctrl: control, EtOH: ethanol-exposed.

**Supplemental Figure 6:**
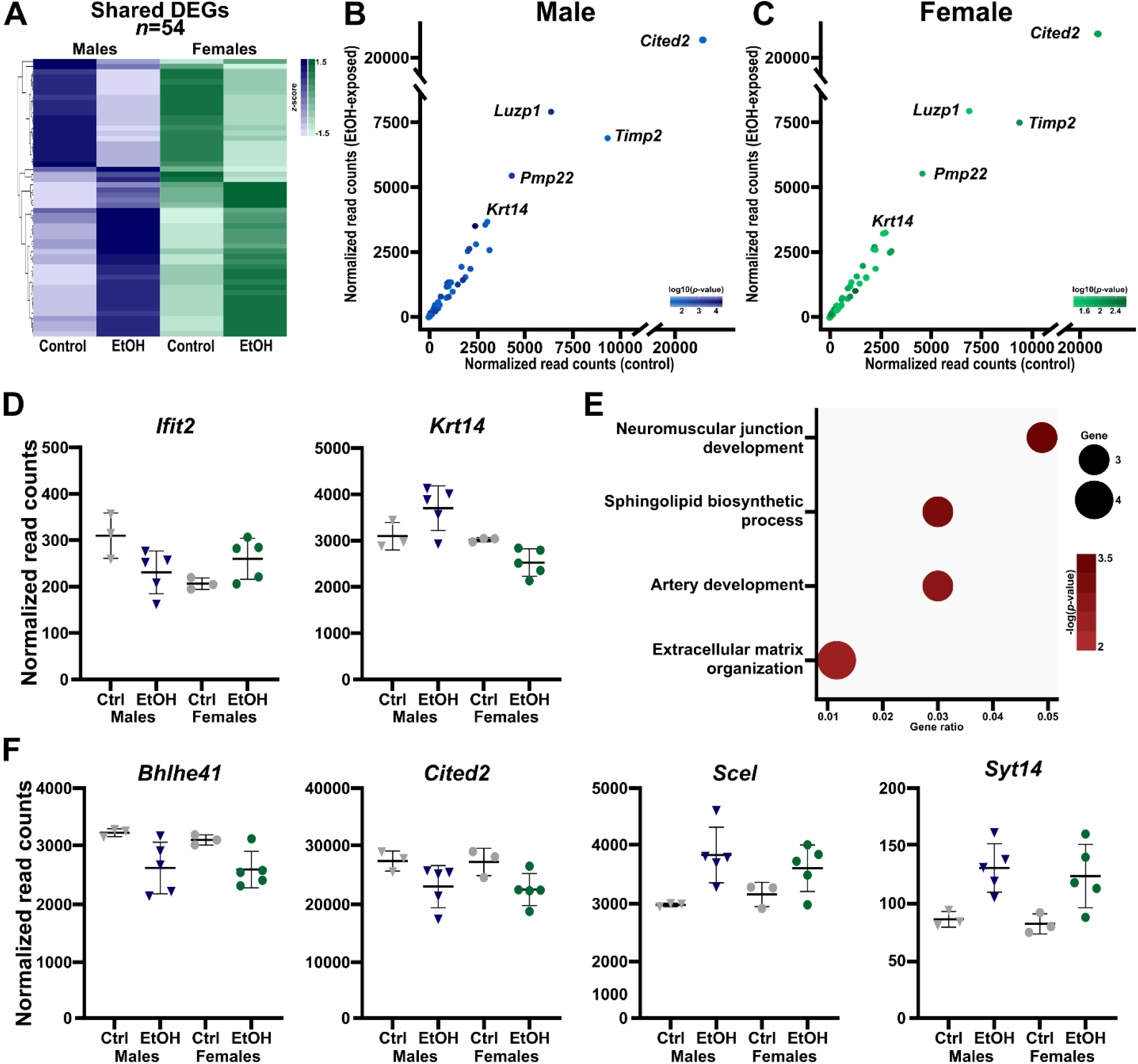
Shared DEGs are similarly altered in male and female ethanol-exposed placentas. **A)** Heatmaps showing expression level (*z*-score) of common differentially expressed genes (DEGs) between male (blue; left) and female (green; right) placentas (54) following ethanol exposure. **B-C)** Scatterplots of the normalized read counts of the 54 common DEGs in **B)** male and **C)** female control and ethanol-exposed placentas. **D)** Normalized read counts of 2/4 genes with opposing expression changes between male and female ethanol-exposed placentas. **E)** Top pathways enriched for common DEGs (54) based on Metascape analysis. **F)** Normalized read counts of genes with altered expression in male and female ethanol-exposed placentas. Data represent the mean ± SD. Ctrl: control, EtOH: ethanol-exposed.

**Supplemental Figure 7:**
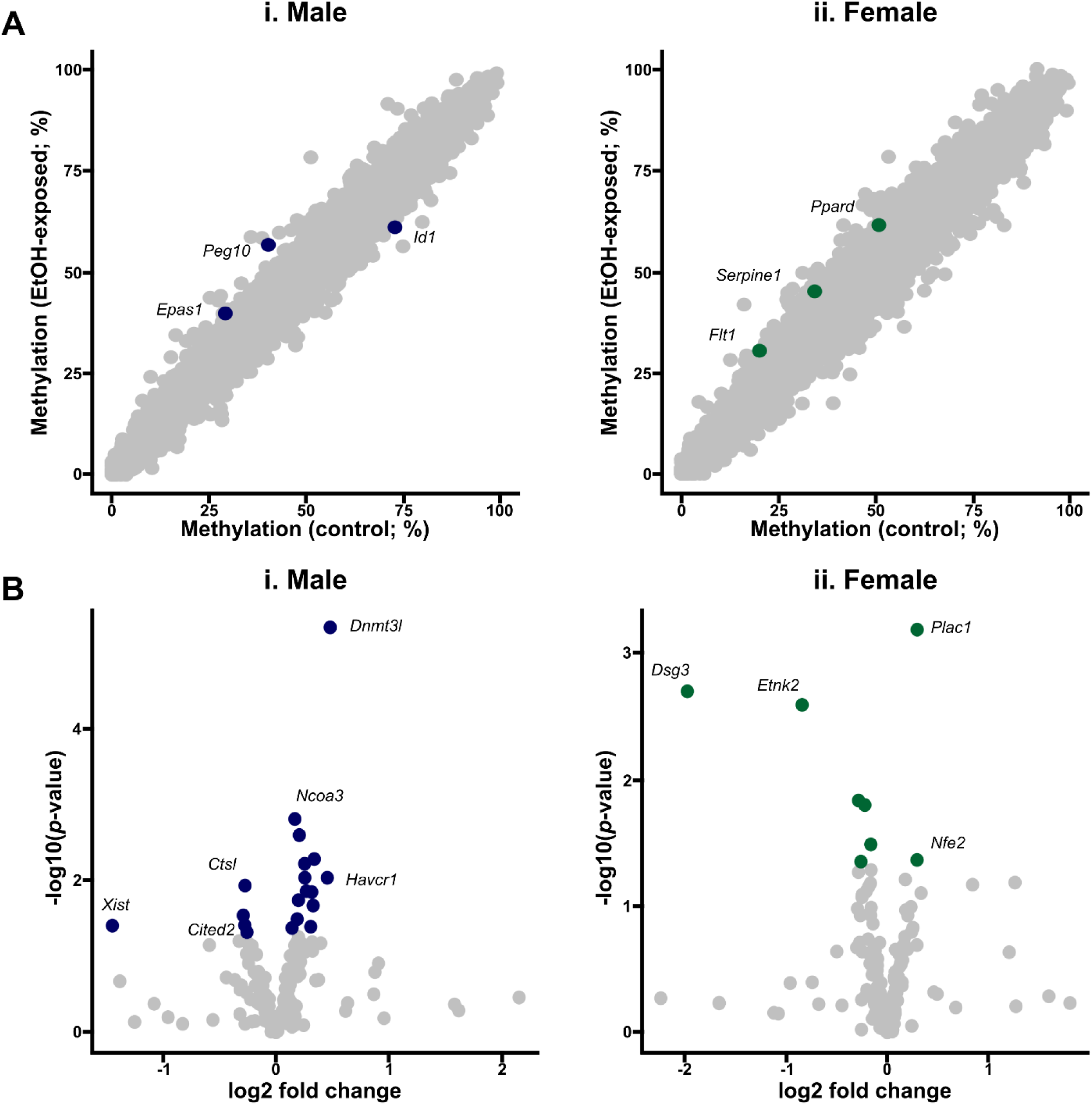
Early preimplantation alcohol exposure selectively alters the epigenetic profiles of placental development genes. **A)** Scatterplot of the tiles located in placental development genes in control and ethanol-exposed **i)** male and **ii)** female placentas. Blue (male) and green (female) dots represent tiles with ≥ 10% methylation changes in ethanol-exposed placentas compared to controls (males, 3; females, 3). **B)** Differential expression analyses of placental development genes between control and ethanol-exposed **i)** male and **ii)** female placentas. Blue (male) and green (female) dots represent genes that were significantly changed (*p*˂0.05) in ethanol-exposed placentas (males, 19; females, 8) compared to controls. Ctrl: control, EtOH: ethanol-exposed.

**Supplemental Figure 8:**
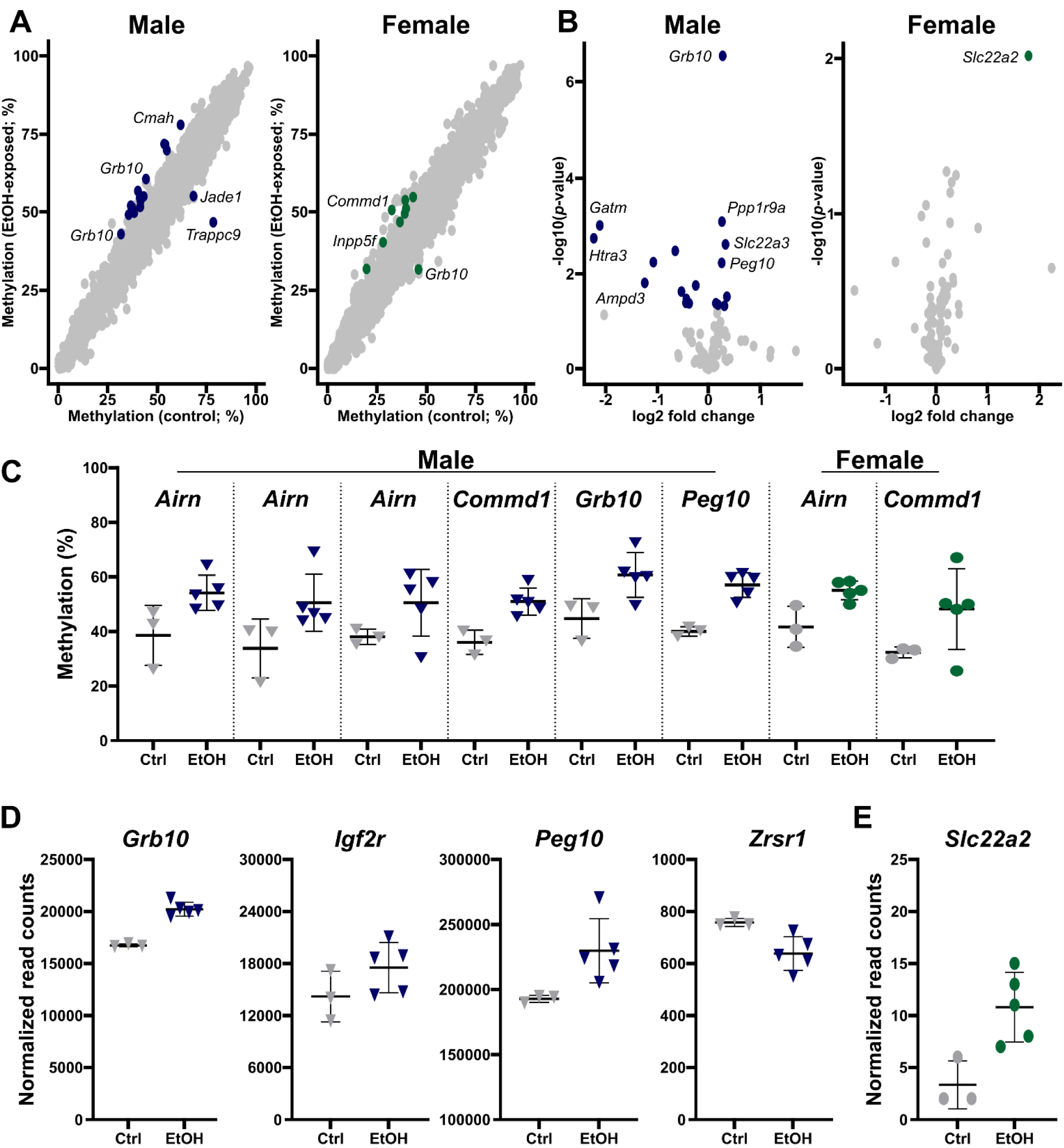
Early preimplantation alcohol exposure alters epigenetic profiles of placental imprinting genes in late-gestation placentas. **A)** Scatterplot of tiles located in placental imprinting genes in control and ethanol-exposed **i)** male and **ii)** female placentas. Blue (male) and green (female) dots represent tiles with ≥ 10% methylation changes in ethanol-exposed placentas (males, 17; females, 8) compared to controls. **B)** Differential expression analysis of placental imprinting genes between control and ethanol-exposed **i)** male and **ii)** female placentas. Blue (male) and green (female) dots represent genes that were significantly changed (*p*˂0.05) in ethanol-exposed placentas (males, 18; females, 1) compared to controls. **C)** CpG methylation levels in control and ethanol-exposed male (left; grey/blue) and female (right; grey/green) placentas for DMRs located in placental imprinting genes. **D-E)** Normalized read counts of placental imprinting genes with altered expression in **D)** male (blue) and **E)** female (green) ethanol-exposed placentas compared to controls. Data are the mean ± SD. Ctrl: control, EtOH: ethanol-exposed.

**Supplemental Figure 9:**
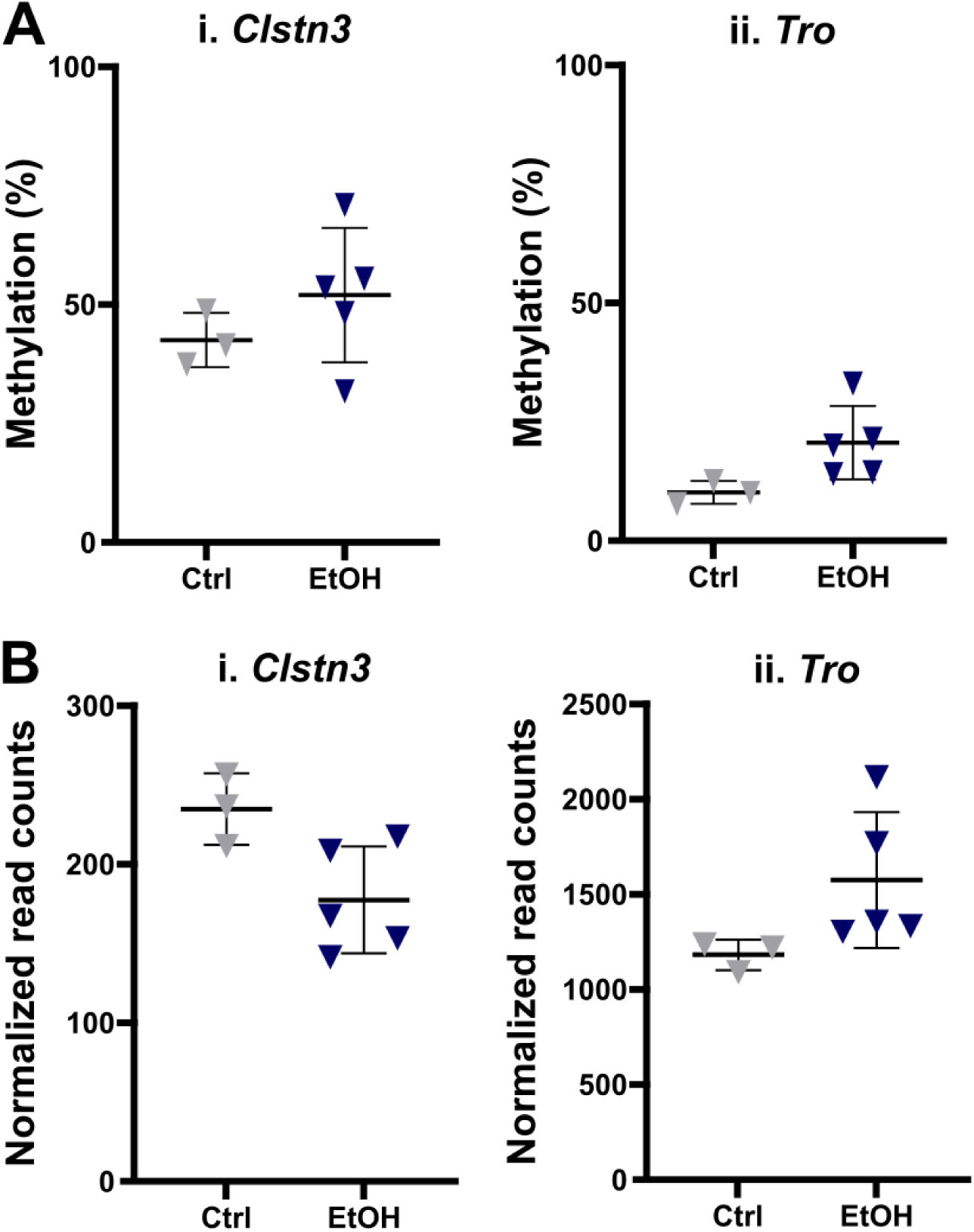
Growth-related genes with altered DNA methylation after ethanol exposure only show expression changes in male placentas. **A)** CpG methylation levels in control and ethanol-exposed male placentas for DMRs located in the DEGs **i)** *Clstn3* and **ii)** *Tro*. **B)** Normalized read counts of **i)** *Clsnt3* and **ii)** *Tro* in control and ethanol-exposed male placentas (*n=*3–5 for each group). All data represent the mean ± SD. Ctrl: control, EtOH: ethanol-exposed.

**Supplemental Figure 10:**
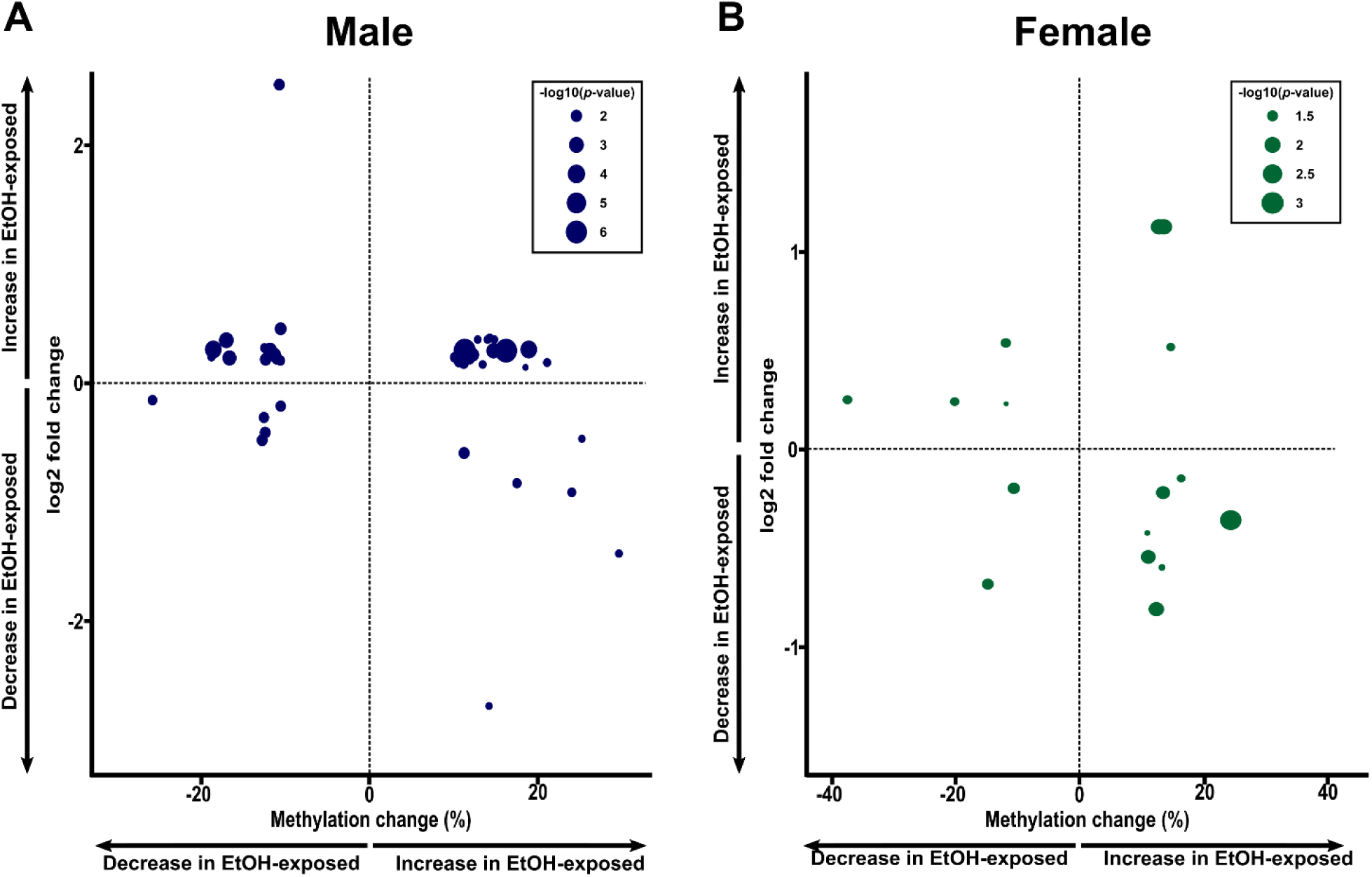
Differential expression is not directly linked to differential DNA methylation in late-gestation placentas following early preimplantation alcohol exposure. DNA methylation differences between control and ethanol-exposed (X-axis) placentas and changes in gene expression (log2 fold change; Y-axis) in regions containing both DMRs and DEGs between control and ethanol-exposed placentas in **A)** males and **B)** females. Ctrl: control, EtOH: ethanol-exposed.

**Supplemental Figure 11:**
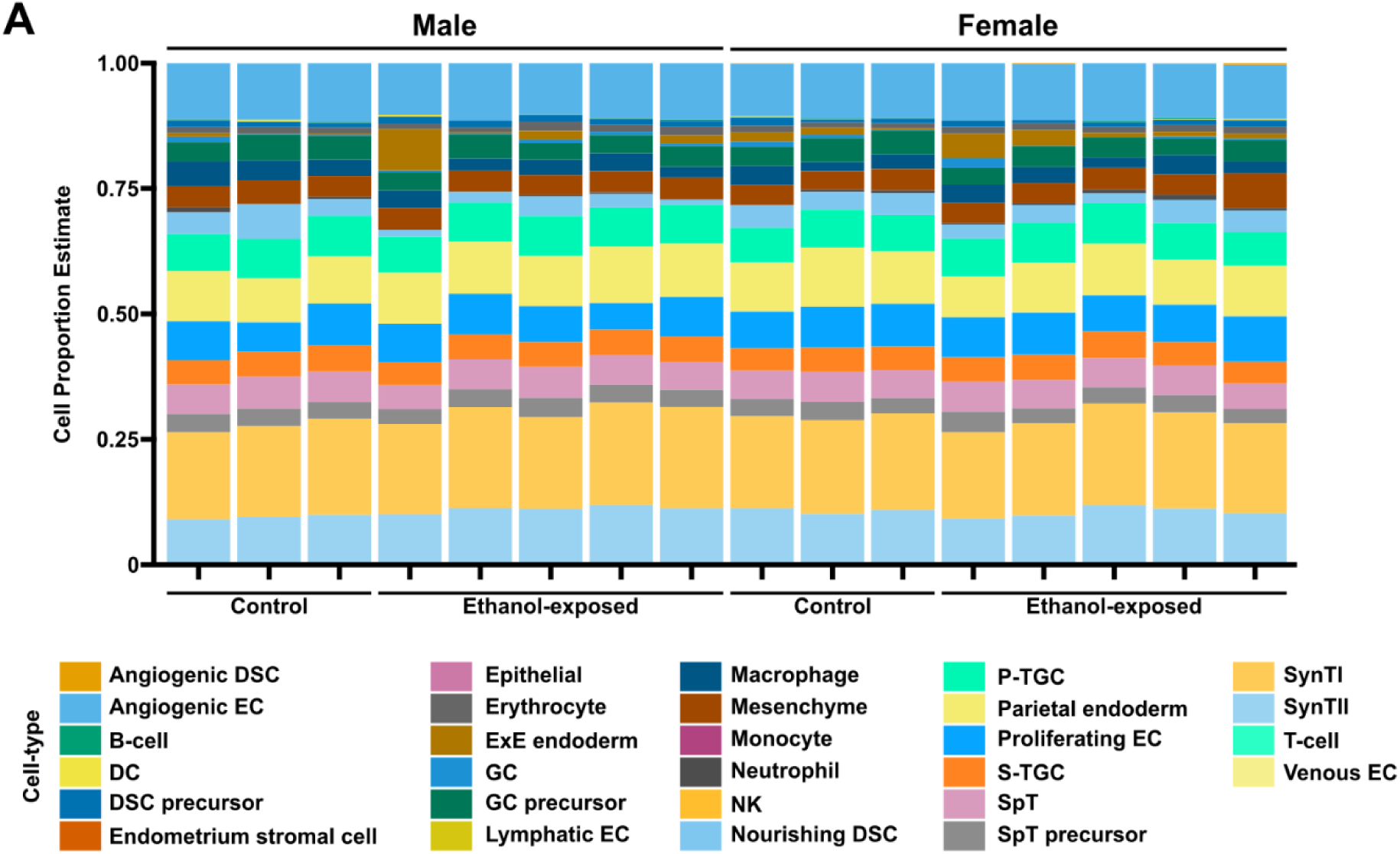
Bulk RNA-seq deconvolution reveals no major distribution shift in E18.5 placental cell subtypes following preimplantation alcohol exposure. Evaluation of the cell composition in bulk RNA-seq of E18.5 mouse placenta by leveraging a deconvolution approach using AutoGeneS (v1.0.4) and the NuSVR model. The final dataset consisted of 9779 cells across 28 cell populations. Figure shows a summary of the cell-type proportion for all samples grouped by conditions (control or ethanol exposed). (Males : Ctrl *n* = 3 ; EtOH *n* = 3. Females Ctrl *n*= 5; EtOH *n* =5). The E18.5 single-nuclei RNA-seq dataset were obtained from (Fu et al., 2024). DC: dendritic cell; DSC: decidual stromal cell; EC: endothelial cell; ExE: extraembryonic; GC: glycogen trophoblast cell; NK: natural killer; P.TGC: parietal trophoblast giant cell; SpT: spongiotrophoblast cell; S.TGC: sinusoid trophoblast giant cell; SynTI: syncytiotrophoblast cell Type I; SynTII: syncytiotrophoblast cell Type II.

**Supplemental Figure 12:**
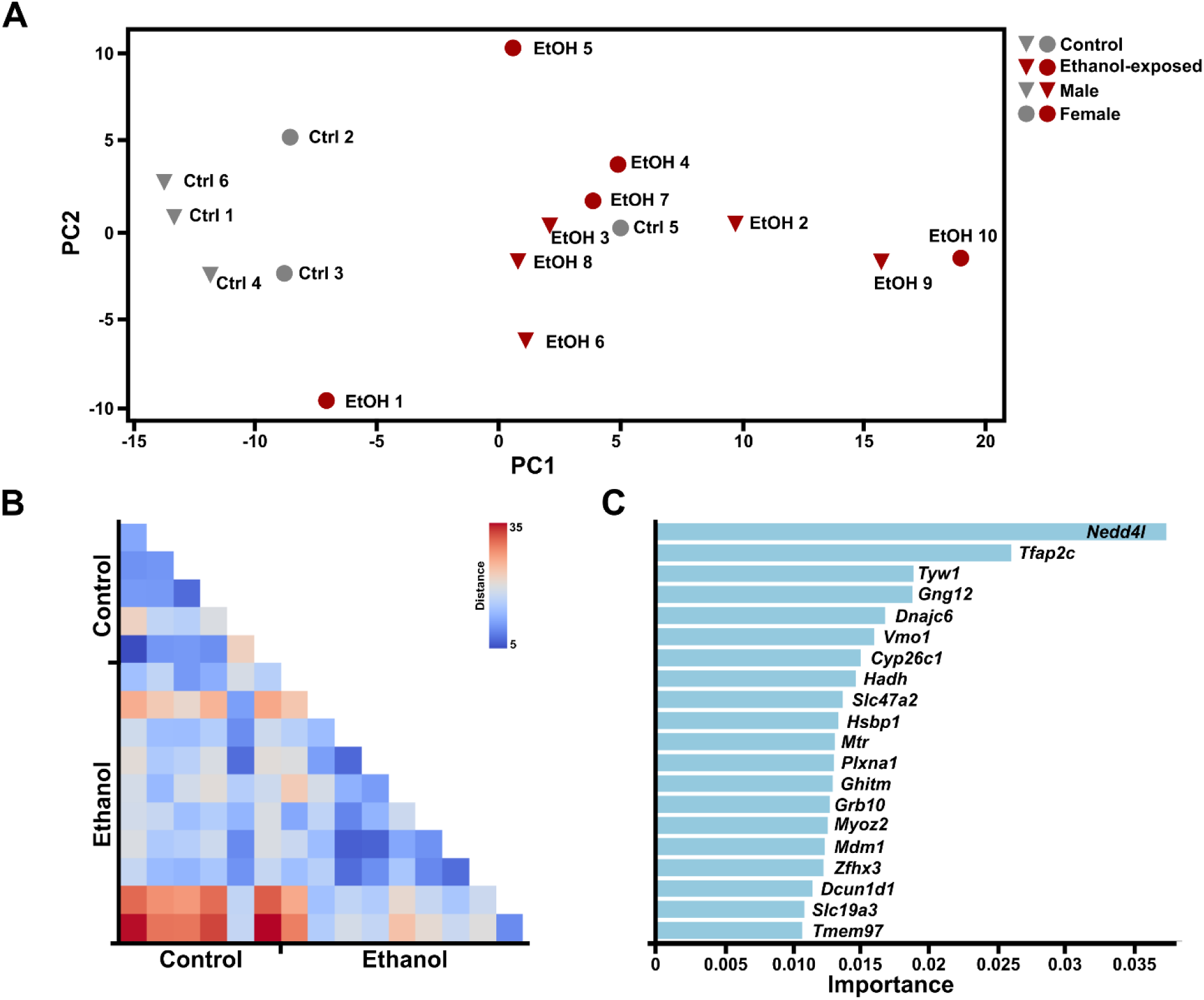
The transcriptomic signature of PAE does not enable PAE detection. **A)** Principal component analysis showing the clustering of control and ethanol-exposed samples based on their gene expression levels in the 20 DEGs making up the signature. **B)** Heatmap of the Euclidean distances of each control and ethanol-exposed sample based on their gene expression levels in the 20 DEGs. **C)** Feature importance plot of the DEGs based on LASSO analysis.

## SUPPLEMENTARY TABLES

**Table S1:** List of Male- specific, female- specific, and shared DMRs. *See Supplementary Data 2*

**Table S2:**
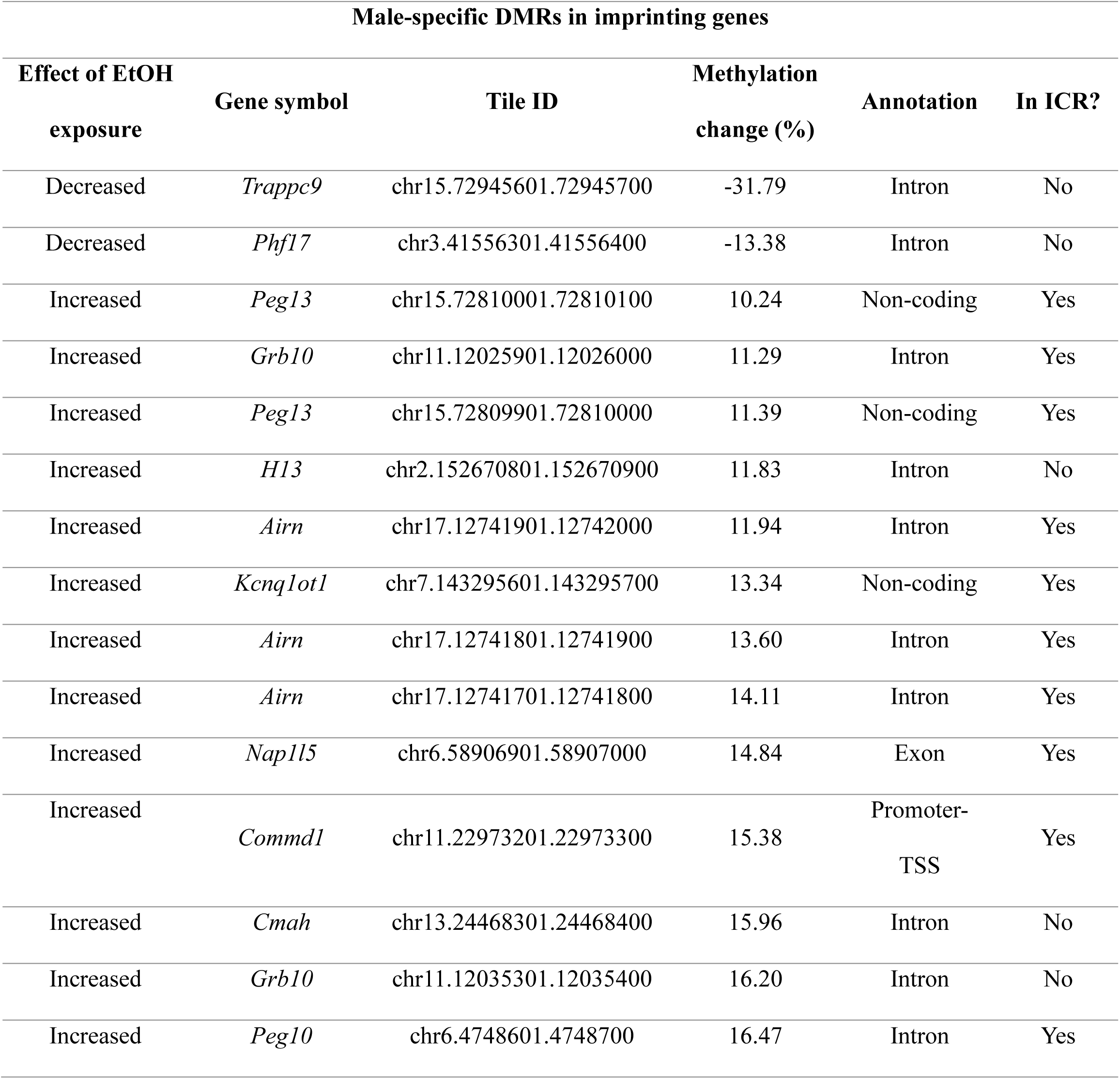
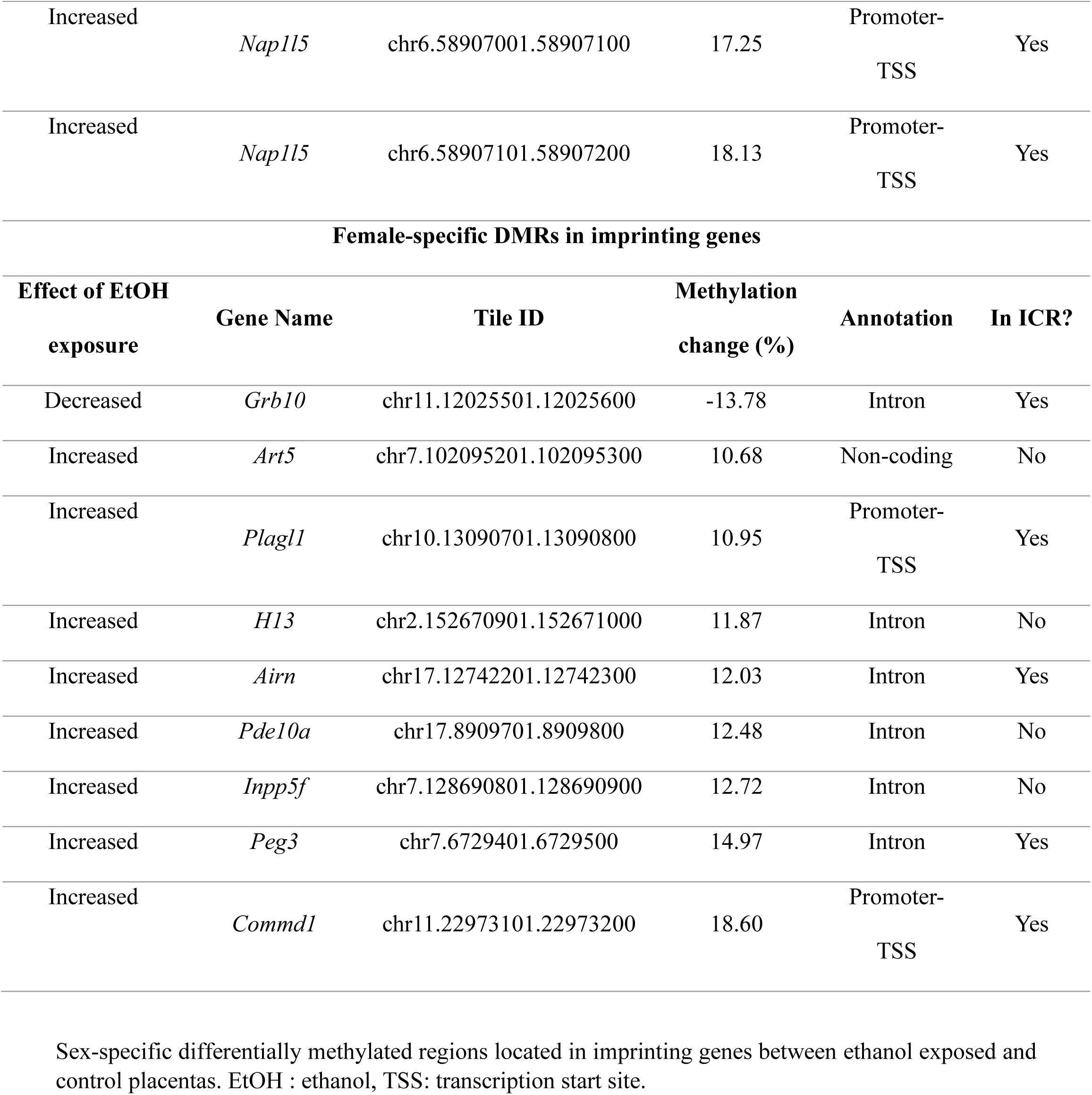
Male- and female-specific DMRs located in imprinting genes.

**Table S3:**
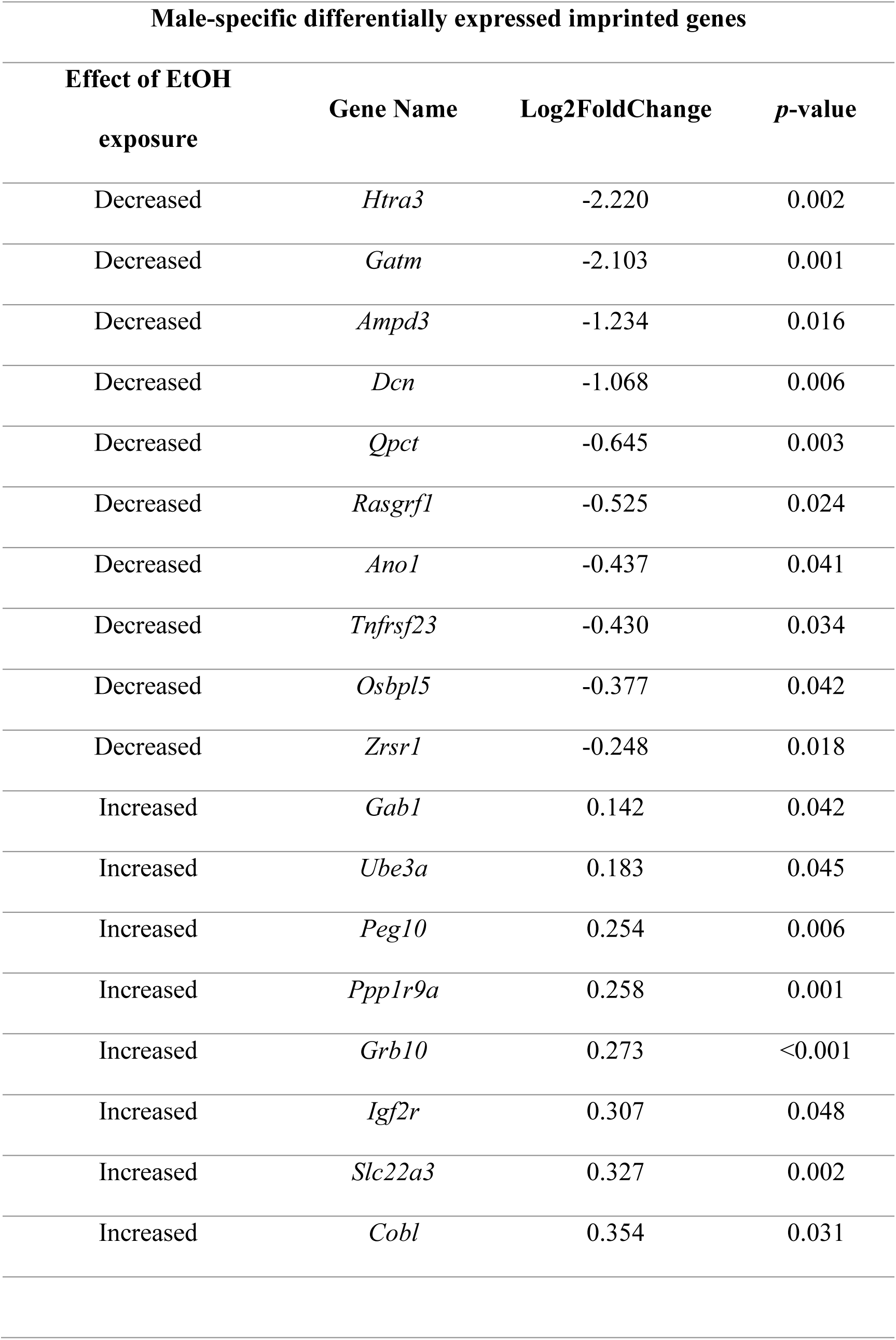
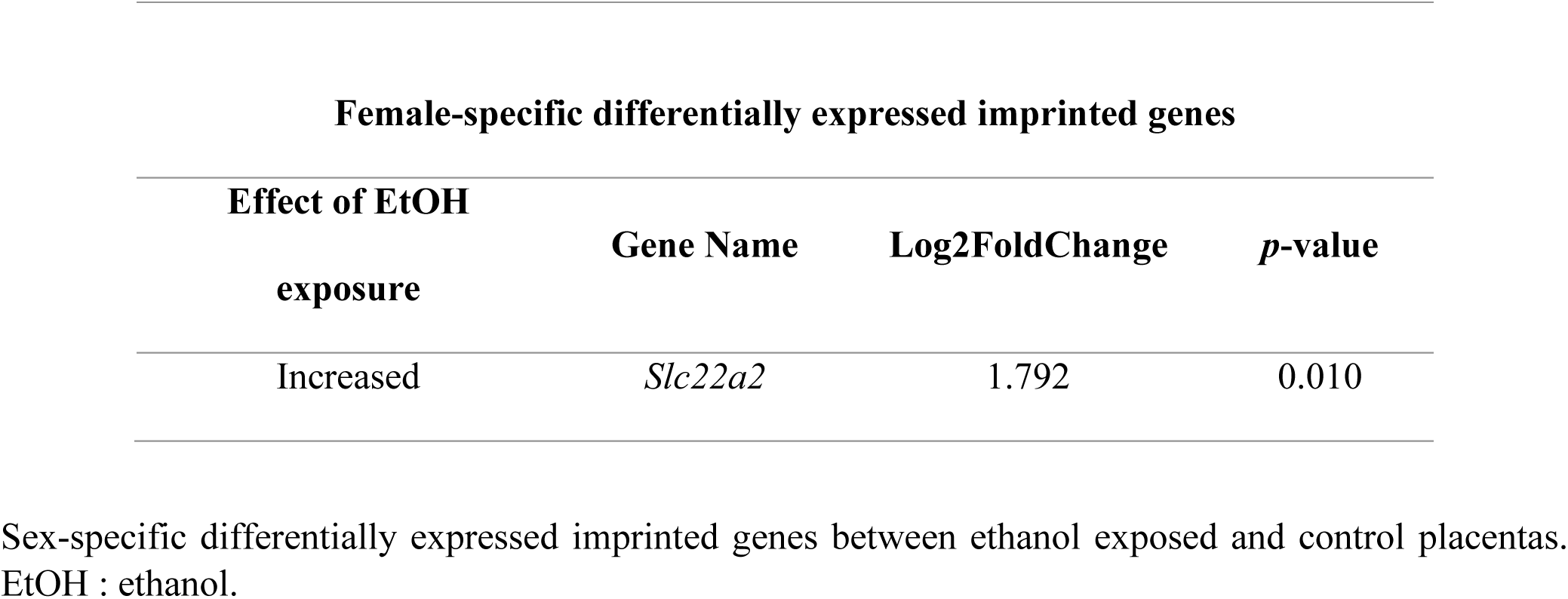
Male- and female-specific differentially expressed imprinted genes.

**Table S4:** Estimated proportion of placental cell composition in male and female between control and ethanol-exposed embryos. *See Supplementary Data 3*

